# Multi-region brain transcriptomes uncover two subtypes of aging individuals with differences in Alzheimer risk and the impact of *APOEε4*

**DOI:** 10.1101/2023.01.25.524961

**Authors:** Annie J. Lee, Yiyi Ma, Lei Yu, Robert J. Dawe, Cristin McCabe, Konstantinos Arfanakis, Richard Mayeux, David A. Bennett, Hans-Ulrich Klein, Philip L. De Jager

**Author notes:** Corresponding author. (P.L.D.).

## Abstract

The heterogeneity of the older population suggests the existence of subsets of individuals which share certain brain molecular features and respond differently to risk factors for Alzheimer’s disease, but this population structure remains poorly defined. Here, we performed an unsupervised clustering of individuals with multi-region brain transcriptomes to assess whether a broader approach, simultaneously considering data from multiple regions involved in cognition would uncover such subsets. We implemented a canonical correlation-based analysis in a Discovery cohort of 459 participants from two longitudinal studies of cognitive aging that have RNA sequence profiles in three brain regions. 690 additional participants that have data in only one or two of these regions were used in the Replication effort. These clustering analyses identified two meta-clusters, MC-1 and MC-2. The two sets of participants differ primarily in their trajectories of cognitive decline, with MC-2 having a delay of 3 years to the median age of incident dementia. This is due, in part, to a greater impact of tau pathology on neuronal chromatin architecture and to broader brain changes including greater loss of white matter integrity in MC-1. Further evidence of biological differences includes a significantly larger impact of *APOEε4* risk on cognitive decline in MC-1. These findings suggest that our proposed population structure captures an aspect of the more distributed molecular state of the aging brain that either enhances the effect of risk factors in MC-1 or of protective effects in MC-2. These observations may inform the design of therapeutic development efforts and of trials as both become increasingly more targeted molecularly.

**One Sentence Summary:** There are two types of aging brains, with one being more vulnerable to *APOEε4* and subsequent neuronal dysfunction and cognitive loss.

## INTRODUCTION

There is extensive heterogeneity among older individuals in terms of their cognitive ability, their trajectory of cognitive decline, and the accumulated burden of aging-related brain pathologies. One study, for example, described over 230 unique combinations of brain pathology found in participants of the Religious Order Study (ROS) and Memory and Aging Project (MAP)(*1*). These two studies were designed to be combined for large-scale analyses, and we refer to them as ROSMAP(*2–4*). Further, in these longitudinal studies of cognitive aging, accounting for six known types of brain pathology still leaves over 50% of the variance in cognitive decline unexplained(*5*), suggesting that there are many as yet unknown biological factors that affect the performance of the aging brain.

Another level of heterogeneity is exemplified by the well-validated genetic risk factor *APOEε4* which has a strong effect on Alzheimer’s disease (AD) susceptibility in individuals of European ancestry as they get older; however, this haplotype’s effect is much weaker in African-American or Latin-x subjects(*6*). Further, its effect also appears to wane as individuals of European ancestry enter advanced age(*7*): there may thus be both inter-individual and temporal heterogeneity in the effects of *APOEε4* on AD risk. More generally, we now have a large set of genetic, environmental, and experiential (such as menopause) risk factors associated with susceptibility to AD or AD-related traits. We do not know whether such risk factors have a modest to moderate effect on AD risk in all individuals or whether they may have a much stronger effect in one subgroup of subjects and have limited or no effect in other subgroups, yielding a modest to moderate effect over the entire population. This question has been challenging to explore because (1) there has not been a clear strategy by which to divide the population of older subjects into subgroups and (2) datasets of high-dimensional data which could be used to address this question empirically were, until recently, too small in size to generate robust results. Two recent efforts took a clustering approach to a different question: resolving subsets of individuals among those who have a diagnosis of AD, using data from one brain region(*8, 9*); these investigators found 2-3 subtypes of individuals with AD who differed by the extent and type of pathology.

Here, we pursue a different question and leverage a rapidly growing collection of transcriptomic (RNA sequencing, RNA-seq) profiles from ROSMAP participants who are all non-demented at study entry. Thus, we are sampling the entire range of states seen in the aging brain (from non-impaired to advanced AD) to define the structure of the older population in a data-driven manner. Further, we consider data from three different brain regions simultaneously, searching for participant subgroups defined by global brain state: we are looking for broader patterns than those which can be found in a single brain region which may relate to the specialized function of that region. Using data derived from ROS and MAP participants, we describe two subtypes of older brains and explore the clinicopathologic and neurobiologic relevance of this structure. Access to transcriptomic profiles from three brain regions – the dorsolateral prefrontal cortex (DLPFC), the posterior cingulate cortex (PCC) and the anterior caudate (AC) – of the same large group of individuals allows simultaneous consideration of all regions, creating a more integrative view of the aging brain. This strategy identified two “meta-clusters”, two groups of aging individuals based on the assessment of multi-region brain transcriptomes, and these empirically defined meta-clusters differ in their rate of cognitive decline, in the integrity of the white matter (not sampled by these transcriptomes), in the biological consequences of Tau pathology and in the impact of the *APOEε4* risk factor. Thus, one of these two groups of participants appears to be more vulnerable to the effect of AD risk factors, while the other is relatively resilient: we therefore define vulnerability as the state within which a risk factor is more likely to result in the accumulation of pathology and symptoms. Finally, we find evidence that these meta-cluster definitions are transportable to other sample collections, addressing the generalizability of our results.

This is the first report of newly enlarged and novel RNA sequence datasets from ROSMAP: DLPFC which includes data from 1,092 participants, AC with 731 participants, and PCC with 661 participants. The AC and PCC data have not yet been analyzed in terms of gene expression changes in relation to AD traits: only ∼10% of these data were used in a recent study focused on RNA editing(*10*). We perform a *de novo* clustering approach to define subsets (meta-clusters 1 and 2) of ROSMAP participants based on the RNA sequence profiles, and, then, we explore the nature of these two subsets in terms of the effect of known risk factors. Thus, the clustering of ROSMAP participants is not based on any prior study or pathophysiologic feature of AD. For the first time, we consider clustering results from 3 regions simultaneously to define our subgroups, unlike other efforts which look at different regions separately and/or consider features that define the presence or stage of AD.

## RESULTS

### Description of the study participants and data

ROS and MAP are two prospective studies of aging that recruit non-demented participants at baseline, follow each participant annually with neurologic and neuropsychological evaluations, and collect brains at the time of death since agreement to brain donation is a prerequisite for participation in the study. Thus, participants capture the diversity of the older population at the time of death, ranging from cognitively non-impaired, to mildly impaired, to dementia in the cognitive spectrum. This allows us to deploy study designs (such as the one used here) that do not rely on categorical variables: we can take advantage of the fact that the ROSMAP participants sample the entire distribution of the older population. The characteristics of ROSMAP participants with transcriptomic data derived from postmortem brains are presented in **Table 1**. After rigorous quality control and our preprocessing pipeline (see Materials and Methods), we retained 19,147 transcripts in 731 participants for the AC, 18,629 transcripts in 1,092 participants for the DLPFC and 19,017 transcripts in 661 participants for the PCC for downstream analyses. A total of 1,149 ROSMAP participants had transcriptomic data in at least one brain region. A subset of 459 participants had RNA-seq profiles in all three brain regions and served as the Discovery cohort (**Fig. 1A**). The remaining 690 participants had RNA-seq profiles in only one or two regions and were used as a Replication cohort. Data generation had been attempted on all available samples, so there is no selection of individuals for data generation in a given region.

**Table 1.** Summary statistics of ROSMAP samples included in the analysis

| Characteristic | AC | DLPFC | PCC | Discovery | Replication |  |  |  | All |
| --- | --- | --- | --- | --- | --- | --- | --- | --- | --- |
|  |  |  |  |  | All | AC | DLPFC | PCC |  |
| Cohort size, *n* | 731 | 1,092 | 661 | 459 | 690 | 272 | 633 | 202 | 1,149 |
| Female, *n* (%) | 478<br>(65%) | 732<br>(67%) | 416<br>(63%) | 284<br>(62%) | 491 (71%) | 194<br>(71%) | 448<br>(71%) | 132<br>(65%) | 775<br>(67%) |
| *APOE*ε4, *n* (%) | 193<br>(27%) | 273<br>(25%) | 165<br>(25%) | 119<br>(26%) | 171 (25%) | 74 (27%) | 154<br>(24%) | 46 (23%) | 290<br>(25%) |
| Age (years) at death, mean (SD) | 89.2 (6) | 89.6 (7) | 89.6 (7) | 89.4 (6) | 89.8 (7) | 89.0 (7) | 89.7 (7) | 90.1 (7) | 89.6 (7) |
| AD dementia, *n* (%) | 237<br>(49%) | 390<br>(52%) | 212<br>(47%) | 140<br>(46%) | 265 (55%) | 97 (54%) | 250<br>(56%) | 72 (48%) | 405<br>(51%) |
| Pathologic AD, *n* (%) | 452<br>(62%) | 688<br>(63%) | 400<br>(61%) | 267<br>(58%) | 460 (67%) | 185<br>(68%) | 421<br>(67%) | 133<br>(66%) | 727<br>(63%) |
| Amyloid, mean (SD) | 1.7 (1) | 1.7 (1) | 1.6 (1) | 1.7 (1) | 1.7 (1) | 1.8 (1) | 1.7 (1) | 1.6 (1) | 1.7 (1) |
| Tau, mean (SD) | 2.2 (1) | 2.3 (1) | 2.1 (1) | 2.1 (1) | 2.4 (1) | 2.4 (1) | 2.4 (1) | 2.2 (1) | 2.3 (1) |
| Cognitive decline, mean (SD) | -0.012<br>(0.1) | -0.015<br>(0.1) | -0.004<br>(0.1) | -0.004<br>(0.1) | -0.021<br>(0.1) | -0.027<br>(0.1) | -0.023<br>(0.1) | -0.004<br>(0.1) | -0.014<br>(0.1) |

**Fig. 1.**
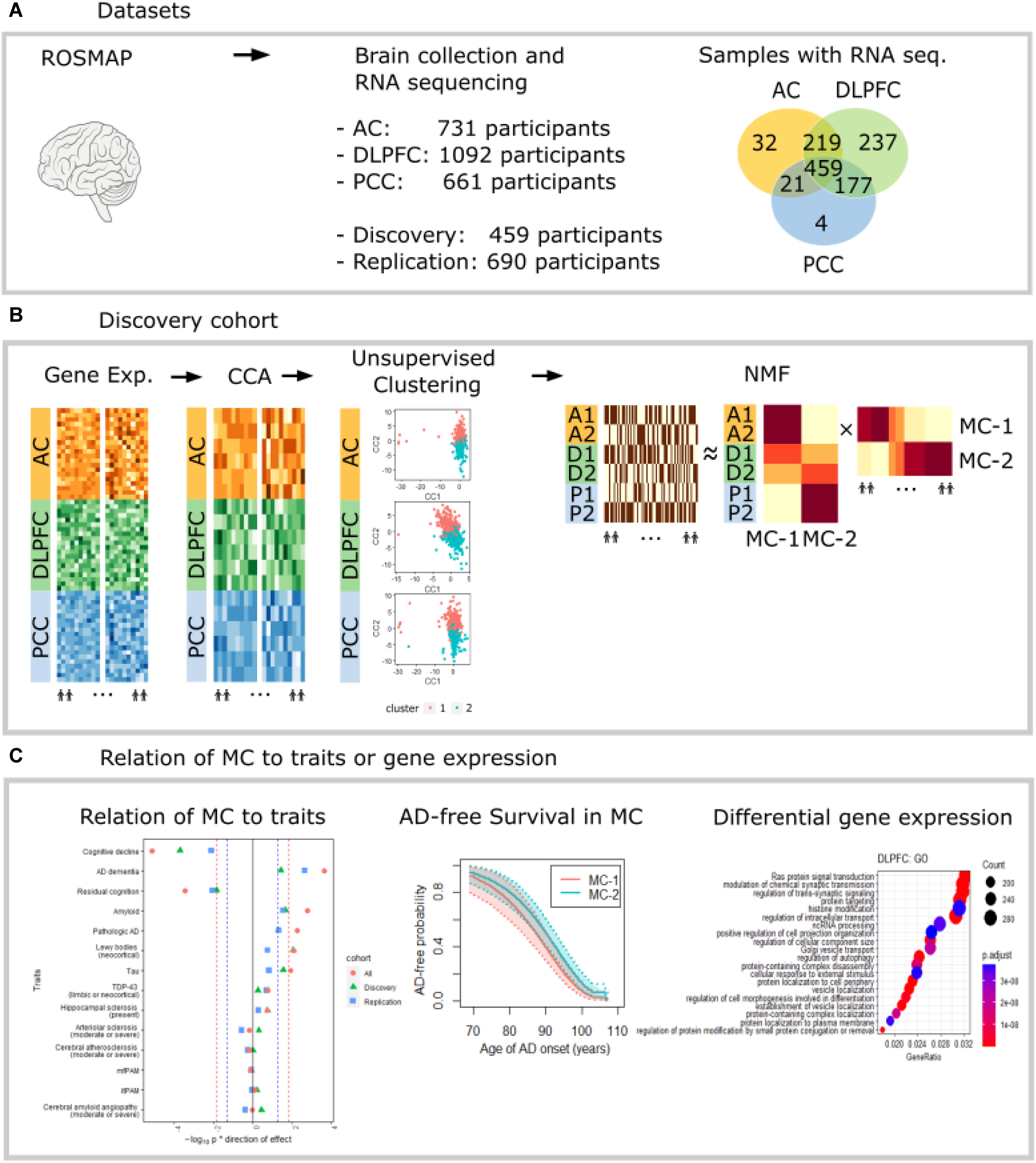
Diagram of the study design. **(A)** The transcriptomic RNA sequencing profiles was accessed from postmortem brains in AC, DLPFC, and PCC regions from ROSMAP. Discovery cohort of 459 participants have RNA sequence profiles in all three brain regions. Replication cohort of 690 participants have RNA sequence profiles in only one or two of these regions. Venn diagram shows the number of participants with transcriptomic data in one or more brain regions. (**B**) For a Discovery cohort of individuals, multiple sparse canonical correlation analysis (CCA) was deployed on the individuals for three regions to identify shared correlation structures across regions. Heatmaps under ‘Gene Exp.’ are created using the 20 randomly selected residuals of the normalized TPM values (y-axis) for the first 10 and last 10 participants (x-axis) in each brain region. Heatmaps under ‘CCA’ are created using the first 6 set of canonical variates (y-axis) returned from deploying sparse multiple CCA on the residuals of the normalized TPM values for the same first 10 and last 10 participants (x-axis) in each brain region. K-means clustering was then applied on the first 6 set of canonical variates for 459 Discovery participants in each region and identified two subgroups of participants in each region as shown in scatter plots under ‘Unsupervised Clustering’. The cluster assignments of each subject in Discovery cohort from three different regions were first combined into a matrix (left heatmap under ‘NMF’). Through non-negative matrix factorization (NMF), we factorized the matrix into a basis matrix representing the projection of original clusters from each region to the meta-clusters (center heatmap under ‘NMF’) and coefficient matrix representing the meta-clusters membership of the individual participants (right heatmap under ‘NMF’). To partition the remaining Replication cohort of individuals, the same signature genes and weights obtained in the Discovery cohort for each region were given to the features/transcripts in the Replication cohort, returning canonical variates for each region. The subgroups of participants were then identified by assigning their canonical variates to the nearest cluster centroid found in the Discovery cohort. In each Discovery and Replication cohort, the resulting clusters of individuals from each region were integrated into meta-clusters of participants through NMF. (**C**) To assess whether clinico-pathologic traits are different between two transcriptomically-defined meta-clusters, a regression model was fitted on Discovery and Replication cohort of individuals separately and evaluated the effect of meta-clusters on the trait. We then conducted a meta-analysis of the two association studies and tested the effect of meta-clusters. We further estimated age at AD onset in two meta-clusters and identified transcripts that were differentially expressed between the two meta-clusters in each of the three regions followed by pathway enrichment analysis. *AC* anterior caudate; *DLPFC* dorsolateral prefrontal cortex; *PCC* posterior cingulate cortex; *CCA* canonical correlation analysis; *NMF* non-negative matrix factorization; *MC* meta-cluster*; Discovery* participants with an RNA-seq profile in all 3 regions; *Replication* participants with an RNA-seq profile in only one or two regions.

### Defining the population structure of aged human brains

We considered statistical approaches to define the structure of the ROSMAP participants based on their molecular profiles derived from transcriptomic data. Unlike prior efforts that evaluated individual regions separately and focused on individuals with AD(*8, 9*), we considered the full spectrum of aged brains (from non-impaired to advanced AD) and explored the hypothesis that there may exist broader patterns in the molecular state of the brain that may not be as apparent if one focuses on a single brain region. Importantly, since cognition is supported by a distributed network of functional centers, we were interested in a more comprehensive view of the brain and thus wanted to simultaneously consider all three regions in our clustering efforts. Brain regions are functionally specialized, and certain types of pathology in the older brain have a predilection for specific regions; since we elected to focus on certain regions involved in cognition, we did not select regions that capture the Braak staging for tau proteinopathy. Selecting such regions would yield an interesting but different study; here, we evaluate the relation of Braak staging to our clusters as part of the downstream analyses.

We elected to use canonical correlation analysis (CCA) (*11*) and subsequent unsupervised k-means clustering in our analytic approach (**Fig. 1B**). CCA finds linear combinations of features (known as canonical variates) that maximize the overall correlation across all regions (see Materials and Methods). Our approach involved deploying CCA in the Discovery cohort of individuals that have transcriptional profiles in all three regions, in order to identify shared correlation structures across the three regions. For each region, the k-means clustering was then applied to the set of canonical variates returned from the CCA to identify participant subgroups (**Fig. S1A-C**). We assumed that the subgroups of participants identified through the clustering would be likely to capture similar sources of variation that are shared across regions. In contrast, principal component analysis (PCA) captures sources of variation within each region independently. We then projected the clusters of participants identified from each region using CCA into a common space through non-negative matrix factorization (NMF)(*12, 13*), yielding participant subsets referred to as “meta-clusters” since they are based on an integrative analysis that considers all 3 regions simultaneously (see **Fig. S2A** and **B**; Materials and Methods). The 459 ROSMAP participants used in this Discovery analysis represented a random subset of the study populations; **Table 1** compares the demographic and clinicopathologic characteristics of the Discovery and Replication subjects.

For all three brain regions, the optimal solution was two clusters of participants, leading to two meta-clusters through NMF. Our approach and parameter selection efforts are described in detail in the Materials and Methods section. In short, to empirically determine the optimal number of clusters in each region, we evaluated all possible models from 2 to 10 clusters, under three clustering methods (k-means clustering, hierarchical clustering and pam), and we evaluated the results using three internal clustering validation measures (connectivity measure(*14*), the Dunn index(*15*), and Silhouette width(*16*)) (see Materials and Methods). For all three regions, two clusters of participants were optimal as the connectivity was minimized and the Dunn index with average silhouette width were maximized under both k-means and hierarchical clustering. Clustering stability was evaluated through clusterwise Jaccard bootstrap mean (CJBM)(*17*) which measures the Jaccard similarities of the original clusters to the most similar clusters in the 1,000 bootstrapped samples. When we considered the k-means and hierarchical clustering methods, for all three regions, two clusters of participants obtained through the k-means clustering were the most stable because both clusters had CJBM above 0.85 (**Fig. S3**).

The two clusters in each region were integrated into two meta-clusters through an NMF multi-region approach (**Fig. S2A** and **B**). The coefficient matrices in **Fig. S2B** show the ROSMAP participants in Discovery cohort, and their meta-cluster membership after projection through NMF. The meta-clusters were stable as the subdivision was evident when visualizing the consensus matrix (**Fig. S2D**). In the Discovery cohort of participants, each individual cluster in the AC and PCC perfectly matched to one of the meta-clusters, as shown in the basis matrix in **Fig. S2A**. We did observe differences in assignments as the DLPFC-only clustering returns clusters that did not align perfectly with the meta-clusters. Nonetheless, the majority of samples in each DLPFC cluster align with the meta-cluster assignment, highlighting one advantage of this approach of looking for a comprehensive view of the brain that minimizes region-specific differences. Regional differences are important and interesting but may obscure larger patterns that influence the function of the distributed cognitive networks and yield symptomatic changes in older individuals.

Having determined our meta-clusters in the Discovery cohort, we took their defining characteristics and used them to partition the remaining subjects which constitute the Replication set of ROSMAP participants that have data in only one or two of the three regions (**Table 1**). Two clusters of participants were identified for each region and integrated into two stable meta-clusters through NMF (**Fig. S1D-F**, **Fig. S2G** and **H**; Materials and Methods); and **Fig. S2H** shows the coefficient matrices for the ROSMAP participants in the Replication cohort. While data generation in each region was done on as many samples as were available at the time RNA was extracted, without any phenotypic selection criteria, participants in the Replication cohort had a higher prevalence of clinical and pathological AD than those in the Discovery cohort. **Table 2** compares the demographic and clinicopathologic characteristics of individuals that have data in one or more brain regions, partitioned by the two meta-clusters. **Table S1** describes the distribution of samples by clusters obtained from each region and meta-clusters in Discovery and Replication cohorts.

**Table 2.** Summary statistics of ROSMAP participants with transcriptomic data in one or more brain regions within MC-1 and MC-2.

| Variable | MC-1 | MC-2 | *p* |
| --- | --- | --- | --- |
| Cohort size, *n* | 529 | 620 |  |
| Female, *n* (%) | 363 (69%) | 412 (66%) | 0.434 |
| *APOE*ε4, *n* (%) | 146 (28%) | 144 (23%) | 0.071 |
| *APOE* |  |  |  |
| ε2/ε2 | 0 (0%) | 5 (1%) | 0.971 |
| ε2/ε3 | 50 (10%) | 92 (15%) | 0.015 |
| ε2/ε4 | 9 (2%) | 14 (2%) | 0.495 |
| ε3/ε4 | 124 (24%) | 125 (20%) | 0.351 |
| ε4/ε4 | 13 (2%) | 5 (1%) | 0.038 |
| Age (years) at death, mean (SD) | 89.6 (6) | 89.6 (7) | 0.919 |
| AD dementia, *n* (%) | 221 (58%) | 184 (45%) | <0.001 |
| Pathologic AD, *n* (%) | 359 (68%) | 368 (59%) | 0.003 |
| Amyloid, mean (SD) | 4.5 (4) | 3.8 (4) | 0.001 |
| Phosphorylated Tau, mean (SD) | 7.8 (9) | 6.4 (8) | 0.006 |
| Cognitive decline, mean (SD) | -0.029 (0.1) | -0.002 (0.1) | <0.001 |
| Braak stage |  |  |  |
| I | 30 (6%) | 45 (7%) | 0.106 |
| II | 44 (8%) | 60 (10%) | 0.079 |
| III | 118 (22%) | 162 (26%) | 0.075 |
| IV | 184 (35%) | 201 (32%) | 0.037 |
| V | 142 (27%) | 134 (22%) | 0.024 |
| VI | 9 (2%) | 7 (1%) | 0.033 |
| CERAD score |  |  |  |
| Definite AD | 183 (35%) | 175 (28%) | 0.001 |
| Probable AD | 195 (37%) | 217 (35%) | 0.020 |
| Possible AD | 44 (8%) | 56 (9%) | 0.323 |

*MC* meta-cluster; *n* sample size; *SD* standard deviation; *AD* Alzheimer's disease; *p* p-value.

### Relation of meta-clusters to clinical and pathologic traits

To evaluate the relevance of our transcriptomically-defined meta-clusters, we assessed whether clinical and pathologic features were different between the two meta-clusters. Since many of these traits were correlated, we imposed a False Discovery Rate (FDR) of 0.05 as our threshold of significance (**Table S2**). We did these analyses in the Discovery cohort of subjects which have transcriptomes from all 3 brain regions. As seen in **Fig. 2A** and **Table S2**, individuals in the Meta Cluster 1 (MC-1) group have a steeper rate of cognitive decline than those participants in MC-2 after adjusting for age and sex in the Discovery sample, and this result is validated in the Replication samples. The meta-analysis (Materials and Methods; **Fig. 2A** and **Table S2**) of both sets of participants provides the final results in which the increased sample size also yields significance for an increased risk of clinical AD dementia at the time of death, consistent with the more rapid decline in cognitive performance seen in MC-1 (**Fig. 2B**). Similarly, our measure of residual cognition(*18*) – a measure of global cognition unexplained by known brain pathologies (i.e., cognitive performance at the last evaluation before death adjusted for all pathologic indices) – is also reduced in MC-1. We also see some difference in a pathologic diagnosis of AD which appears to be driven by higher β-amyloid load in MC-1 (**Fig. 2A** and **Table S2**), with the burden of tau pathology being only marginally higher in MC-1 (p_FDR_=0.023). Neocortical Lewy bodies were also higher in MC-1 (p_FDR_=0.019). By contrast, the burden of several other pathologies contributing to cognitive decline –cerebrovascular pathologies, TDP-43, and hippocampal sclerosis – are not significantly different between the two meta-clusters. To address whether the difference (p_FDR_ = 0.0001) in the rate of cognitive decline between MC-1 and MC-2 is driven by differences in the burden of AD pathology between the two groups, we repeated the analysis and adjusted for the burden of β-amyloid and tau pathology, the two defining pathologic characteristics of AD. The result remains significant (p_FDR_ = 1.04 × 10^−4^) (**Table S3**), and therefore the difference in cognitive decline comes from another, as yet uncharacterized mechanism. This is also illustrated in **Fig. 2C** in which the meta-cluster assignment remains associated with rate of cognitive decline when adjusted for all the neuropathologies and covariates. The meta-clusters did not segregate individuals with and without dementia (**Fig. 2D**): there is a large proportion of demented participants among both MC-1 and MC-2. There was an increased frequency of dementia in MC-1, but this was not a defining characteristic of that group. Results are similar if one considers a pathologic diagnosis of AD using the Reagan criteria (**Fig. 2E**). When we stratified the participants by AD status to explore the data further in secondary analyses, MC-1 had a steeper rate of cognitive decline than MC-2 among participants with pathological AD (p_FDR_=0.004), after adjusting for age and sex. The result remains significant when we additionally adjusted for the burden of β-amyloid and tau pathology (**Table S5**).

**Fig. 2.**
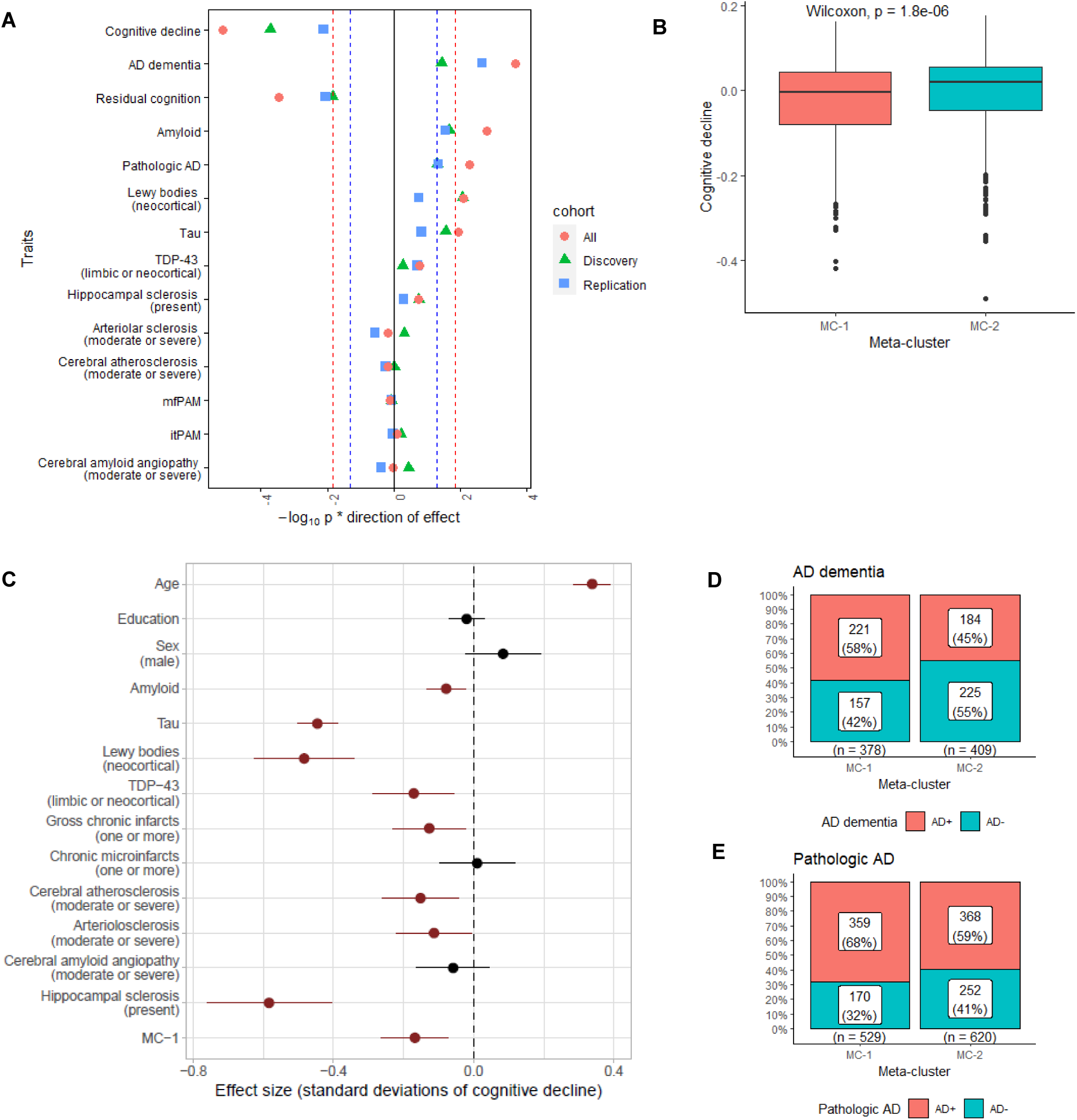
Clinico-pathologic traits that are different between transcriptomically-defined meta-clusters of participants. (**A**) Association of each trait between MC-1 and MC-2 after adjusting for age and sex in Discovery (green triangle) and Replication (blue square) cohorts, and meta-analysis across the Discovery and Replication cohorts, called “All” (red circle). -log10(*p*), weighted by direction of effect, indicates the strength of evidence for the association. The red dotted lines indicate FDR thresholds of 0.05 using the tests in the table, and the blue dotted lines indicate uncorrected thresholds of *p* = 0.05. All *p*-values are two-sided. (**B**) Boxplot of the slope of cognitive decline in MC-1 and MC-2 for 1,149 ROSMAP participants with transcriptomic data in one or more brain regions. (**C**) Factors which independently influence slope of cognitive decline when adjusted for all the other neuropathologies and covariates. A linear regression model was fitted on the 1,149 participants with their meta-clusters assignments from the Discovery and Replication cohorts. The red dots indicate significant association with p-value < 0.05. Line segments represent the 95% confidence intervals. (**D, E**) Bar chart of diagnosis of AD and pathological diagnosis of AD in MC-1 and MC-2 from the 1,149 participants.

### Differences in other factors related to cognitive decline

To further explore what could be driving some of the difference in cognitive decline between MC-1 and MC-2, we evaluated a summary measure of epigenomic perturbation caused by Tau pathology that we derived from genome-wide histone 3 lysine 9 acetylation (H3K9ac) data(*19*). This score reflects a summary measure of large-scale chromatin alterations and captures a proximal functional consequence of tau proteinopathy: the extent of chromatin relaxation in affected neurons. Although we cannot fully capture the state of chromatin with a single chromatin mark, our carefully selected mark does capture important changes in chromatin structure that relate to Tau pathology. There may be other chromatin changes that we have not yet appreciated, but the H3K9Ac-related changes are large, widespread in the genome, and were validated *in vitro*(*19*). This epigenomic tau score was associated with the rate of cognitive decline (b=-0.0004, p=0.0006) in ROSMAP participants. Interestingly, the burden of tau pathology, which is based on the amount of pathologic phosphorylated tau aggregates that have accumulated in the brain(*20*) is only marginally different between the two meta-clusters. Yet, the epigenomic tau score was greater among MC-1 than MC-2 (b=16.3, p=1.04 × 10^−6^), and the difference in epigenomic score remained highly significant when adjusted for the burden of tau pathology (**Table S4**). This suggests that some of the molecular events downstream of tau that contribute to cognitive decline are poorly captured by neuropathological counts of pathologically phosphorylated tau fibrils.

Most of the outcome measures that we evaluated so far relate to cortical function and neuropathologies. Since we aimed to capture a comprehensive view of the brain with our multi-region modeling approach, we also investigated whether the meta-cluster assignment captured aging-related alterations in white matter function. In ROSMAP, post-mortem magnetic resonance imaging (MRI) was used to evaluate loss of brain tissue integrity that relates to cognitive decline. One of the previously characterized MRI measures is the transverse relaxation rate R2, an important tissue characteristic that is the multiplicative inverse of T2 and is thought to reflect the ratio of free to bound water molecules in the tissue. We had previously defined a region of frontal white matter where R2 explains some of the variance in cognitive decline, after accounting for known neuropathologies(*21*). This region of frontal white matter has a difference in R2 level between the two meta-clusters (**Fig. S4** and **Table S4**). This result suggests that our meta-cluster classification, based on three gray matter RNA-seq profiles, also captures an element of disruption in white matter integrity. Moreover, the difference in the accelerated cognitive decline seen in MC-1 remained significant when adjusted for the white matter R2 (**Table S3**), so it is only one aspect of the different state of MC-1 brains. When we stratified the participants by AD status to explore the data further in secondary analyses, MC-1 had a steeper rate of cognitive decline than MC-2 among participants with pathological AD (p_FDR_=0.005) (**Table S5**), after adjusting for age, sex, amyloid, tau, and R2. Moreover, the average R2 level was lower in MC-1 compared to MC-2 among participants with pathological AD (p_FDR_=0.012) (**Table S5**). We note that these analyses are limited by the smaller sample sizes of the partitioned subsets of participants.

In terms of genetic risk factors leading to cognitive decline, we found no difference in the frequency of the *APOEε4* haplotype (**Table 2** and **S4**) between the two meta-clusters. We do note that there are some differences in haplotype combinations that were not significant after accounting for the testing of multiple hypotheses (**Table S4**): MC-1 participants were more likely to be homozygous for *APOEε4* and less likely to have the *APOEε2/ε3* combination when compared to MC-2. Further, we repurposed polygenic scores for AD that had previously been reported to be associated with a variety of clinical and molecular outcomes in these participants(*22*), but we found that MC-1 and MC-2 have a similar genetic propensity for AD using this summary measure (**Table S4**). When we evaluated common AD susceptibility variants(*23*) in **Table S6**, only the Clusterin (*CLU*) variant displayed nominal significance in allele frequency between MC-1 and MC-2 (p=0.007), adjusting for age and sex as well as technical factors. It was no longer significant after correcting for testing multiple hypotheses (p_FDR_=0.129). MC-1 is more likely to have copies of risk allele in *CLU* compared to MC-2. However, this difference in allele frequencies is small, and, when we add this SNP as a covariate, MC-1 remains strongly associated with cognitive decline (p_FDR_= 4.9× 10^−5^) (**Table S3**), suggesting that this variant is not a major driver of the two clusters of participants. This suggests that the meta-cluster structure may not be related to susceptibility but rather as a modulator of responses to risk factors and pathologic events.

To summarize these results and offer a representation of the relative importance of our meta-cluster assignment, we present the proportion of variance in cognitive decline explained by all pathologies and the additional variance explained by the meta-clusters (**Fig. 3**); the meta-clusters explain another 1.1% of variance (p=0.003), beyond the known neuropathologic factors and the measure of frontal white matter integrity. For comparison, using a similar model, *APOEε4* did not significantly explain the variance of this trait, consistent with prior reports that the haplotype’s effect is primarily mediated by effects on pathology(*24*). For comparison, neocortical lewy body burden explain another 2.4% of variance (p=1.6× 10^−6^). Thus, the effect of our meta-cluster structure on cognition is meaningful and distinct from known pathology.

**Fig. 3.**
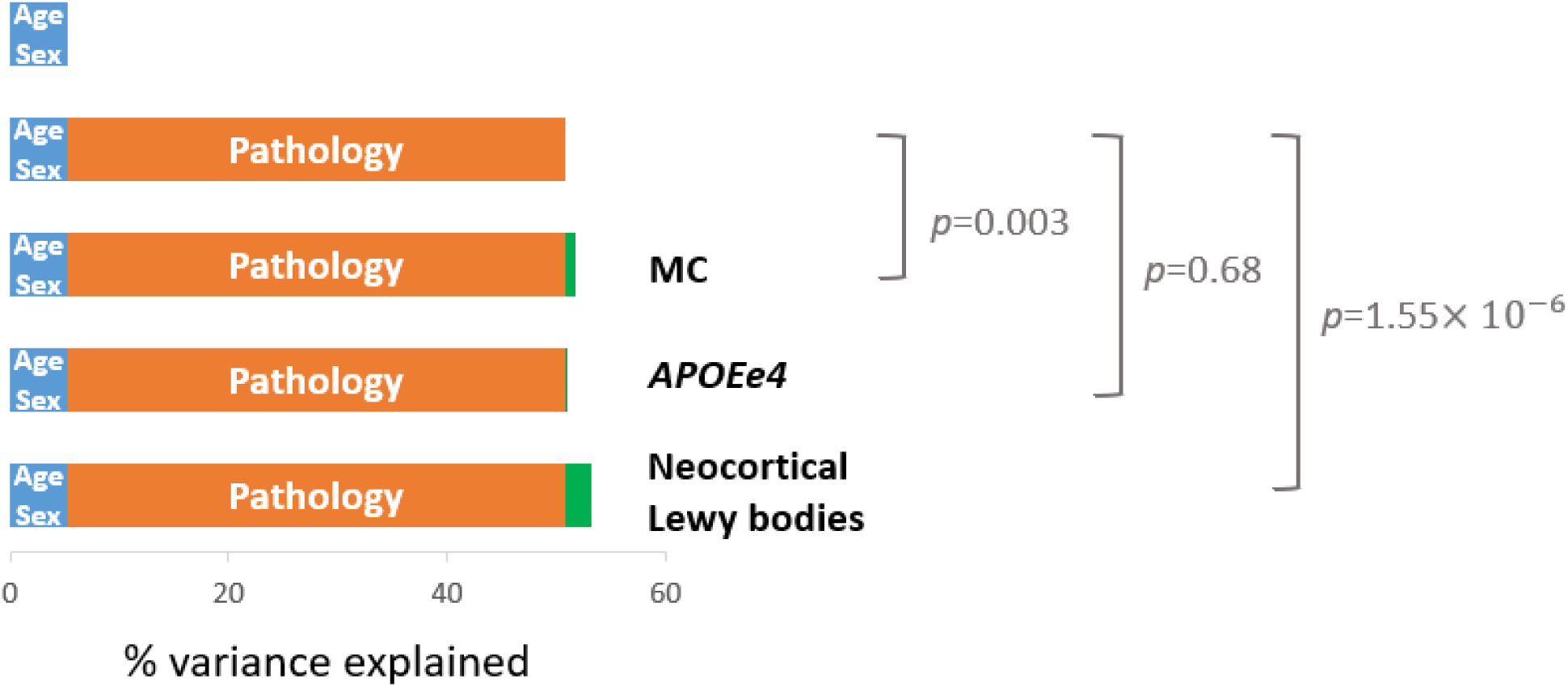
Percentage of variance in slope of cognitive decline explained by each model. In blue, we denote the proportion explained by age and sex alone. The pathological traits that we considered are: amyloid, phosphorylated tau, tdp-43 (limbic or neocortical), cerebral atherosclerosis (moderate or severe), cerebral amyloid angiopathy (moderate or severe), arteriolosclerosis (moderate or severe), hippocampal sclerosis, and R2. Their joint effect is shown in orange. We then performed a linear regression model, fitted for slope of cognitive decline with their meta-clusters assignments on the 503 of 1,149 participants who have all the complete traits, which explained a small (1%) but significant proportion of unexplained variance in cognitive decline (shown in green). On the other hand, *APOEε4* does not explain any more variance, consistent with prior work that suggest that its effect works primarily through AD pathology. For comparison, neocortical lewy body burden explain another 2.4% of variance, beyond the known neuropathologic factors and the measure of frontal white matter integrity.

### Pathway analysis of differentially expressed genes

To annotate the group of genes that define the two meta-clusters, we next identified those transcripts that were differentially expressed between the two meta-clusters in each of the three regions through differential gene expression followed by pathway enrichment analysis using the functional Kyoto Encyclopedia of Genes and Genomes (KEGG) Pathway database(*25*) with FDR correction (FDR<0.05). The estimated proportion of neurons(*26, 27*) was included as a covariate to the regression model in order to account for the effects of possible changes in neuronal cell populations.

The 12,723 genes that were differentially expressed between MC-1 and MC-2 in DLPFC displayed enrichment for 87 KEGG functional pathways (**Fig. 4A**). Gene sets related to several different neurodegenerative diseases were enriched (including AD, amyotrophic lateral sclerosis, and Parkinson’s disease), suggesting that the meta-cluster may capture some shared element of CNS vulnerability or resilience. There was also less pronounced enrichment for the endocytosis pathway that has been implicated in AD. Results from the AC and PCC are very similar (**Fig. 4A**). We did not see APOE-related pathways emerge in these analyses.

**Fig. 4.**
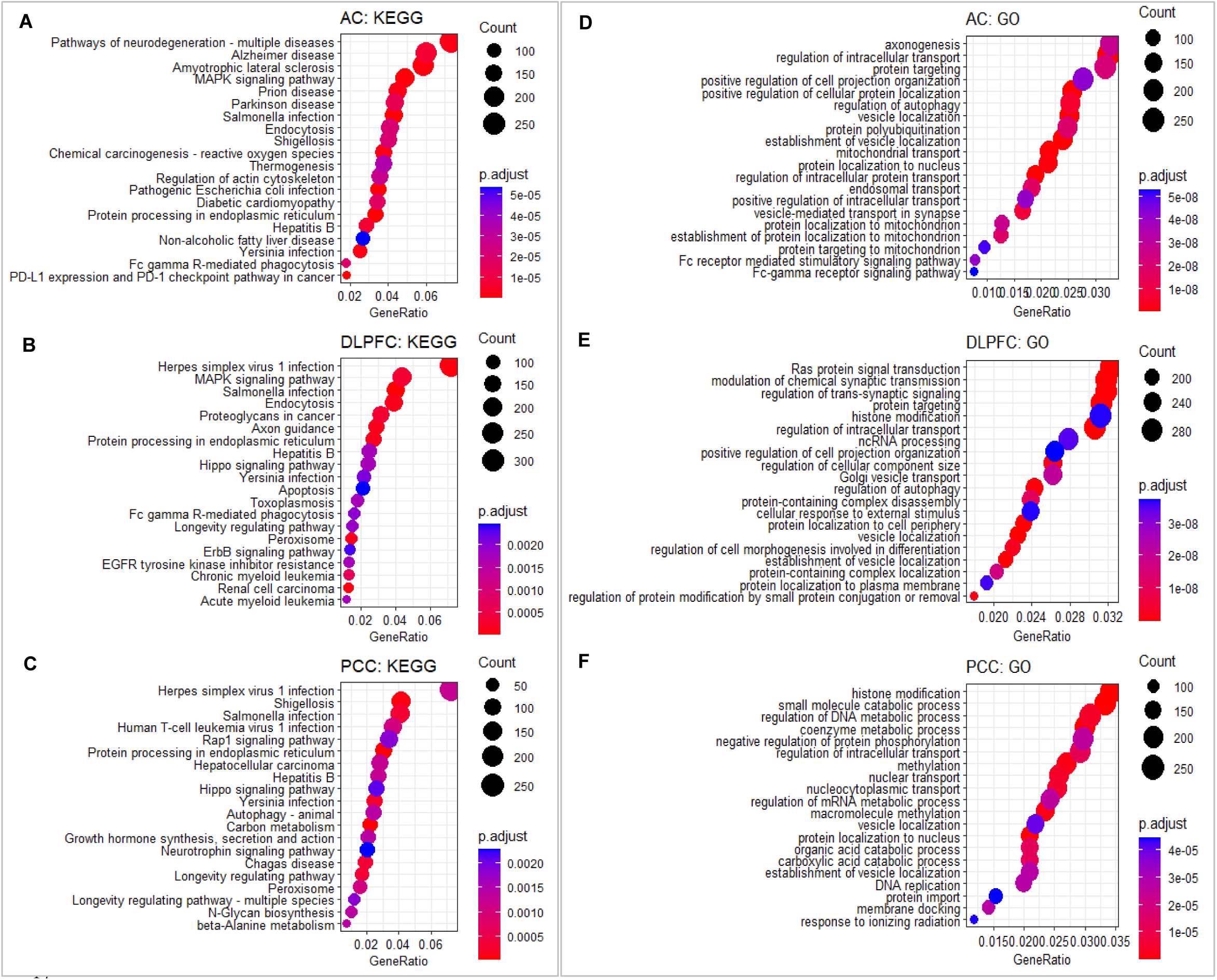
KEGG pathways analysis and gene ontology enrichment analysis in AC, DLPFC, and PCC. (**A-C**) Top 20 enriched KEGG pathways of the differentially expressed genes comparing MC-1 to MC-2 in three brain regions. (**D-F**) Top 20 enriched gene ontology biological processes of the differentially expressed genes comparing MC-1 to MC-2 in three brain regions (FDR adjusted p-value < 0.05).

We also conducted gene ontology (GO) enrichment analysis of the differentially expressed genes comparing MC-1 to MC-2 in each region, and results are shown in **Fig. 4B**. Here, protein targeting and neuronal signatures related to synaptic transmission and trans-synaptic signaling are prominent, suggesting that neurons may be driving part of this difference, although the proportion of neurons in each sample was accounted for.

### Transcriptome-derived cluster structure propagates into the proteome

While we are limited to a smaller subset of participants and one region (DLPFC) with proteomic data in ROSMAP, we nonetheless evaluated whether the differences in RNA expression that define the meta-clusters propagate to the protein level. In **Fig. 5A**, we compare the proteins and RNA expression levels in the same (DLPFC) region. The detected proteins were distributed along the RNA expression spectrum with a skew towards the more abundant RNA expression; when we compare the fold-change between MC-1 and MC-2 of a gene to that of the protein it encodes, we find a correlation of 0.21 (𝑝𝑝 < 2.2 × 10^−16^), illustrating that the differences in molecular state between the two meta-clusters are present at the protein level and consistent with the RNA level. GO enrichment analysis of the detected proteins revealed some of the same pathways that we found in the RNA expression, such as the cellular protein complex disassembly (*p* = 7.60× 10^−11^) and protein-containing complex disassembly (*p* = 1.28× 10^−8^) (**Fig. 5B**).

**Fig. 5.**
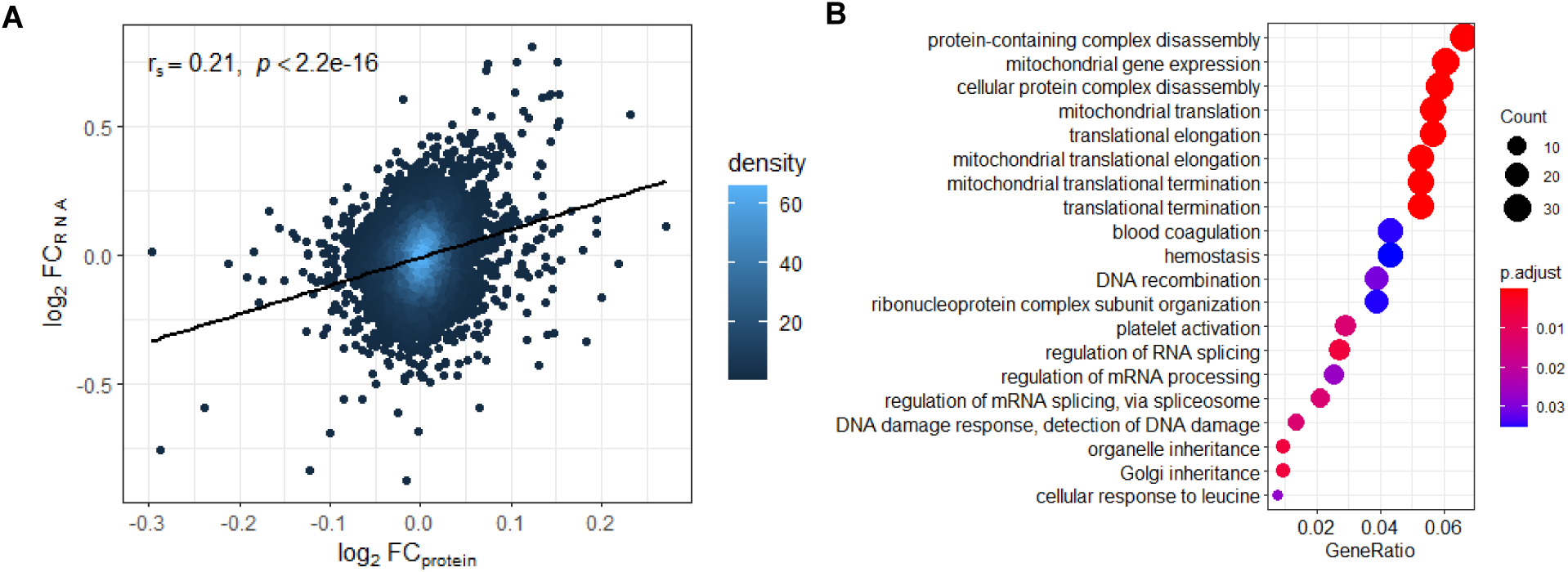
Differential expression analysis of proteomic data in DLPFC. (**A**) Scatter plot of the fold change between the MC-1 and MC-2 subsets for proteins (x-axis) and RNA (y-axis) expression levels in DLPFC. Spearman correlation (𝑐𝑐_𝑠𝑠_) was used. (**B**) Top 20 enriched gene ontology biological processes of the differentially expressed proteins in DLPFC (FDR adjusted p-value < 0.05) between MC-1 and MC-2 were considered.

### Differences in cell type proportions

To address the possible role of changes in cell type proportions in the meta-cluster definition, we evaluated each cell type using signatures derived from purified cell populations(*19, 27*), and we observed that the inferred proportion (from the bulk DLPFC tissue profile) of almost all cell types was different between the two meta-clusters (**Table S4**). There appears to be a broad effect on the cellular composition of the CNS parenchyma that differentiates MC-1 from MC-2. To address whether this is a late manifestation of AD due to greater neuronal or other cellular loss in individuals with AD, we repeated the analysis in the subset of ROSMAP participants who died without cognitive impairment (no dementia and no mild cognitive impairment), and we found that all cell types (except oligodendrocytes) were also different in these cognitively non-impaired individuals. Thus, a lower inferred proportion of neurons and higher proportion of other cell types is an early feature of MC-1 subjects, being present prior to cognitive impairment. While this analysis cannot tell us whether the observed differences in cell types are the product of brain development and/or early, asymptomatic neurodegeneration, the cell population differences are clearly not simply a reflection of the terminal phase of AD.

To confirm these results, we accessed targeted proteomic data in which peptides encoded by the synaptosome-associated protein 25 (*SNAP25*) gene were measured in a set of 1,092 ROSMAP participants in the DLPFC, many of which are included in the RNA based analyses. *SNAP25* is an important synaptic protein and can therefore offer a proxy of synaptic health. It is also reduced in MC-1, consistent with the reduced inferred proportion of neurons (**Table S4**). The result remained significant when we adjusted for the proportion of neurons estimated from the transcriptome (*p* = 6.38× 10^−4^). This suggests that the difference in *SNAP25* may not be largely driven by neuronal loss. When we adjusted for *SNAP25*, we find that proportion neurons are still reduced in MC-1 (*p*=5.06 × 10^−28^) (**Table S4**). This suggests that *SNAP25* levels indicate synapse loss, a phenotype that is related to but different loss of the neuronal soma.

For microglia, we also tested a different set of genes: the HuMi-Aged gene set that we have previously defined as being enriched in microglia isolated from aged ROSMAP brains(*28*). This signature is also increased in MC-1, and the difference is much more significant than the inferred microglial proportion measure (**Table S4**). While all of these signatures derived from bulk tissue homogenates imperfectly captured cell type proportions, the microglia-related results suggested that, even if the proportion of microglia may be increased to some degree in MC-1 because of diminished neuronal proportions, the transcriptional profile of these microglia appear to be more profoundly altered: MC-1 microglia appear to be more “Aged” than MC-2 microglia.

### Heterogeneity in the effect of known AD genetic risk factors in the two subgroups

To investigate the mechanism for the difference in clinical phenotypes between the two groups, we investigated the possibility that an AD risk factor could have a different effect size on the rate of cognitive decline in MC-1 vs. MC-2, which would lead to an excess of symptoms in MC-1. We tested the difference through meta-analysis of Discovery and Replication cohorts (Materials and Methods). We first investigated the *APOEε4* haplotype given its unique role in AD and found that there is a difference (*p*=0.003) in the effect of *APOEε4* between meta-clusters (**Fig. 6** and **Table S7**). Specifically, *APOEε4* was associated with a more rapid cognitive decline in MC-1 individuals relative to MC-2. At least part of this difference in effect may be attributable to enhanced accumulation of tau pathology, as we also see modest interaction of meta-cluster with *APOEε4* in relation to this trait (*p*=0.037) (**Table S7**). However, when we account for the burden of β-amyloid and tau pathology, the interaction of meta-clusters with *APOEε4*, though attenuated, remained significant. Thus, we find that an aspect of the molecular differences between MC-1 & MC-2 appears to be differences in the magnitude of the effect of the *APOEε4* haplotype, with MC-2 individuals having a diminished impact of *APOEε4*. When we evaluated all *APOE* haplotype combinations, the effect of *APOE* on cognitive decline was not significantly different between meta-clusters (p=0.190) (**Table S7**). Among participants with pathological AD, the effect of *APOEε4* on cognitive decline was different in MC-1 and MC-2 (p=0.005) (**Table S8**). To assess whether this interaction is driven by differences in aged microglia, we adjusted the analysis for the HuMi-Aged geneset, but the interaction term remains significant (**Table S7**). Similarly, we adjusted for *SNAP25* expression to assess whether the interaction may relate to neuronal proportion and/or synaptic health; the resulting interaction effect is somewhat attenuated but remains significant (**Table S7**).

**Fig. 6.**
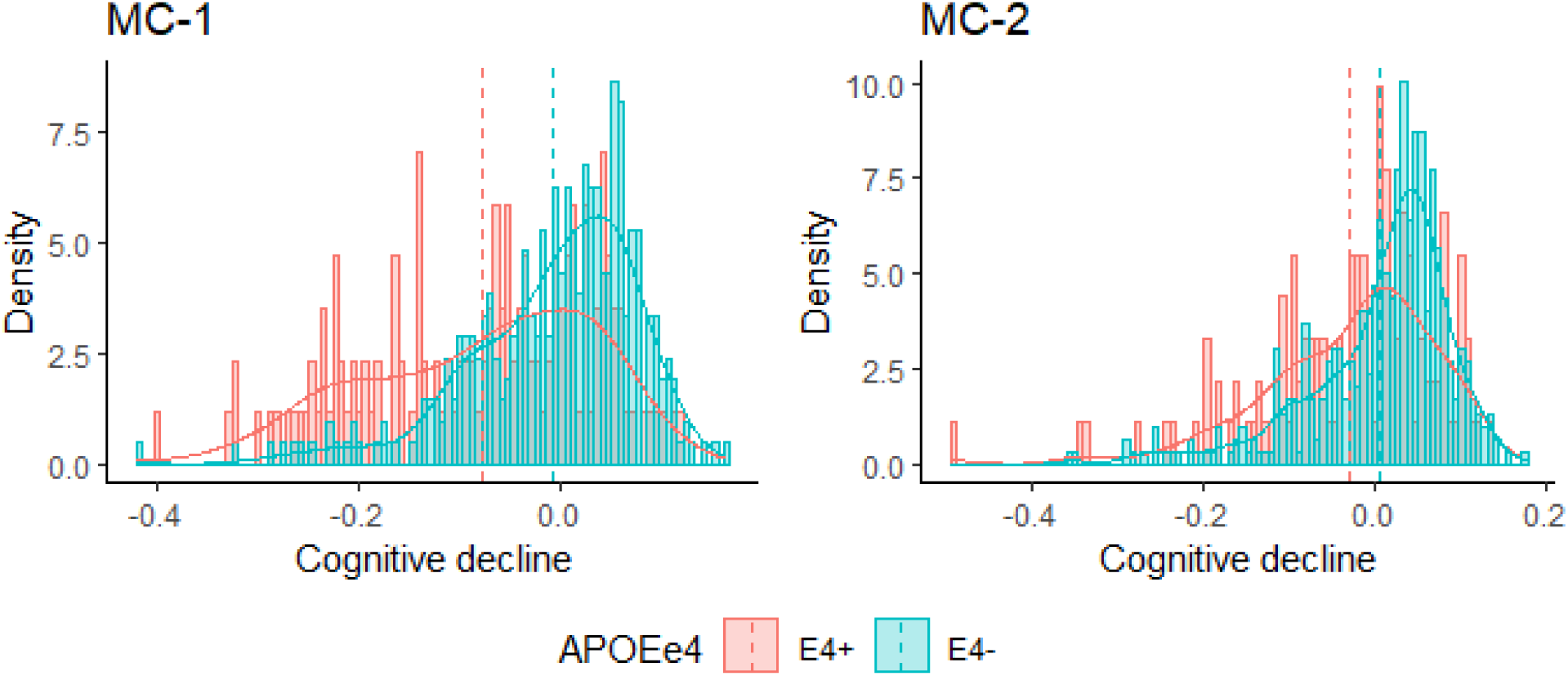
Histogram of slope of cognitive decline by *APOEε4* within MC-1 and MC-2. A significant shift in the distribution of slopes of cognitive decline is seen for *APOEε4* carriers between the MC-1 and MC-2 subsets.

Secondarily, we evaluated other associations with AD-related traits for evidence of interaction, and there was suggestive evidence (*p*=0.024) that MC-1 individuals accumulate more tau proteinopathy per unit of β-amyloid pathology (**Table S7**). This observation will need further validation. We did not find evidence for a significant difference in the effect of the R2 measure of white matter integrity or the other pathologies.

Finally, we recently reported the existence of an epigenomic factor related to microglial activation that enhances the effect of *APOEε4*: the Epigenomic Factor of Activated Microglia (EFAM)(*29*). We therefore hypothesized that MC-2 subjects may have lower EFAM levels, explaining the reduced effect of *APOEε4.* While the level of the EFAM measure itself was not different (*p*=0.087) between MC-1 and MC-2 in the frontal cortex of the 456 ROSMAP participants that also have DNA methylation data (**Table S4**), higher EFAM was associated with more rapid cognitive decline only in MC-1, yielding some evidence for interaction (*p*=0.029) (**Table S7**). However, EFAM does not explain the difference in *APOEε4* effect: when considering both EFAM and *APOEε4* in the same model, we find that the interaction between *APOEε4* and meta-cluster on cognitive decline remained significant (**Table S7**).

### Clinical relevance of meta-cluster assignment

To assess the potential clinical impact of belonging to one of the two meta-clusters of aging brains, we evaluated age at AD onset for MC-1 and MC-2 by conducting a survival analysis using the onset of AD dementia as the primary outcome. As seen in **Fig. 7A**, MC-1 individuals tend to develop dementia earlier than MC-2 individuals. Specifically, if we look at the median age of dementia onset, belonging to the MC-2 meta-cluster leads to a 3 years delay in the median age of dementia onset. Using a different approach, the estimated cumulative risk of Alzheimer’s dementia onset by age 75 in MC-1 was 16.3% (95% CI: 5.2-30.1%), whereas the cumulative risk of Alzheimer’s dementia to age 75 for MC-2 was 11.6% (95% CI: 4.2-21.4%) (**Fig. 7B** and **Table S9**). We adjusted for sex and education. Thus, there is a meaningful difference in clinical outcome that arises from belonging to MC-2.

**Fig. 7.**
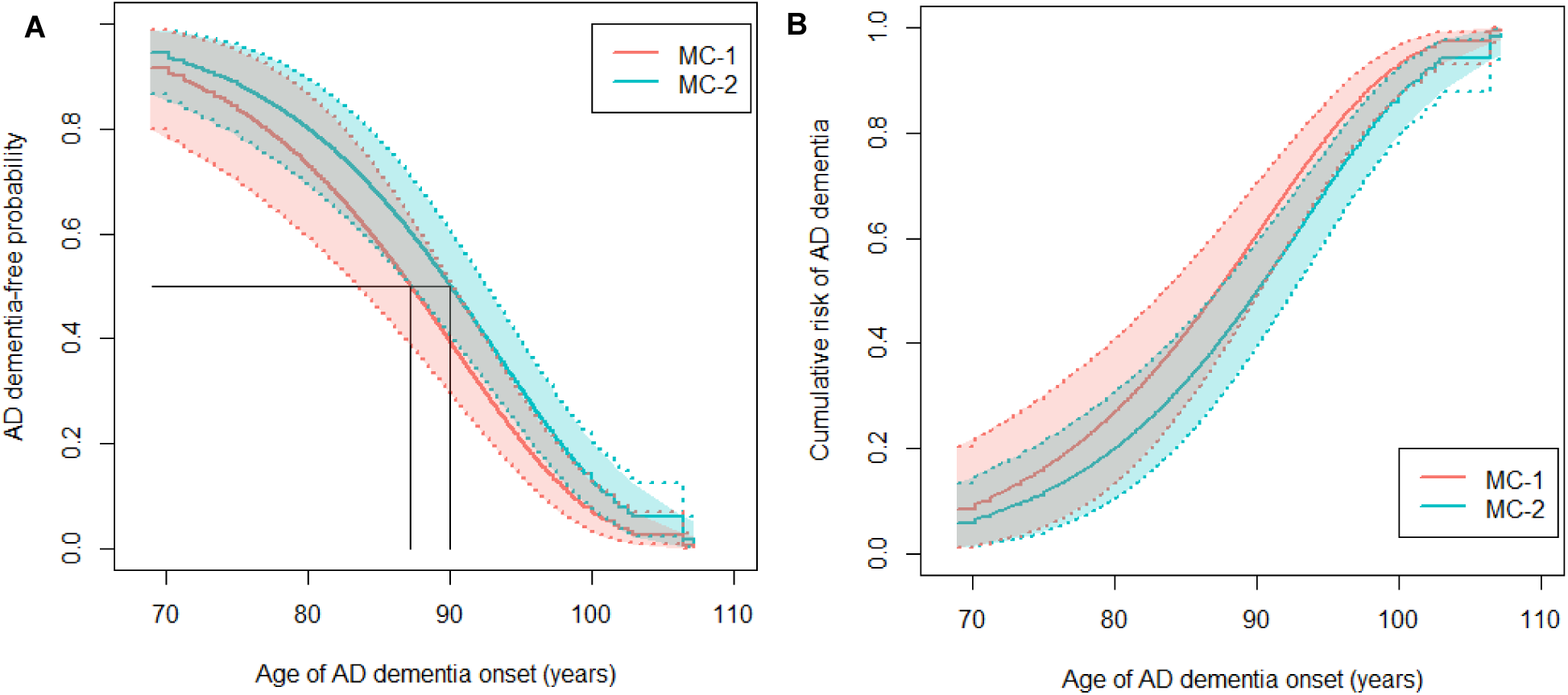
Estimated age at AD dementia onset in MC-1 and MC-2. (**A**) Estimated AD dementia-free survival curves and (**B**) cumulative risk of AD dementia onset in MC-1 (red solid line) and MC-2 (blue solid line) with their 95% confidence intervals (dashed lines), adjusting for sex and education.

### Validation

To assess the generalizability of our observation, we accessed neocortical RNAseq datasets available on the AD Knowledge Portal; we note that the study designs that yielded these data is very different from that of the ROSMAP cohorts, as samples were collected through traditional brain banks and samples selected to fulfill certain categorical criteria. Specifically, we retained 186 participants with data from the inferior frontal gyrus from the Mount Sinai Brain Bank (MSBB)(*30*) and 262 participants with temporal cortex data from the Mayo Clinic (Mayo)(*31*) collection. Using the signature genes and weights from the DLPFC which were used to partition the ROSMAP replication dataset, we obtained canonical variates for both the MSBB and Mayo samples. The subgroups of participants were then identified by deploying the k-means clustering to the canonical variates, leading to two clusters of participants for each cohort. We then assessed whether pathologic diagnosis of AD or interaction of the effect of *APOEε4* on the pathological AD (the relevant traits available in MSBB and Mayo) were different between the two meta-clusters in each cohort. The meta-analysis from ROSMAP, MSBB, and Mayo provides the final results.

Despite a much smaller sample size and not having the most important cognition-related traits in these external replication samples, we found evidence of replication in both Mayo and MSBB (**Fig. S6**). Specifically, there was a difference in pathological AD between MC-1 and MC-2 in Mayo (b=1.565, p=1.83× 10^−5^), and, while this analysis was not significant in MSBB, the estimated effect was consistent with our hypothesis. As seen in the Forest plot (**Fig. S6A**), a meta-analysis using all three cohorts was significant (b=0.237, p=0.045) (**Table S10**). MSBB and Mayo datasets are both much smaller than the ROSMAP dataset and only have pathologic outcomes which have a weaker effect in ROSMAP than the cognition-related traits; nonetheless, we see encouraging evidence that our meta-cluster structure may be generalizable. We note that Mayo has the youngest brains and that this may explain, in part the observed results as MC-1 participants are likely to be over-sampled in their AD group among individuals who are deceased at a younger age, given our observation that MC-1 participants are diagnosed with AD about 3 years earlier, on average. Finally, we did not find evidence for replication of an interaction of the effect of *APOEε4* on a pathologic diagnosis of AD with MC-1 in Mayo and MSBB; however, this result is difficult to interpret given the much reduced size of these two sample collections (**Fig. S6B** and **Table S11**).

## DISCUSSION

We explored the structure of the older population based on the molecular state of their brain at the time of death. Empirically defined subtypes of individuals were determined and found to be related to cognitive decline. While earlier attempts have been made to define subtypes of individuals with AD in other sets of brain data and earlier versions of ROSMAP data(*9, 32*), we improved on prior work in two important ways: (1) we had a much larger sample size, with up to 1,149 individuals with brain transcriptomes and (2) we had a large number of ROSMAP participants (*n*=459) which had transcriptomes from three distinct brain regions involved in cognition (the AC, DLPFC and PCC). Thus, we had greater statistical power to resolve the presence of small subsets of subjects, if present, and we could deploy a different approach which considered the three regions simultaneously. The latter approach may provide a better, integrative view of the state of an individual’s brain that overcomes technical factors and biological idiosyncrasies of the data from each region. We also considered the full range of aged ROSMAP participants and not just the subset with AD, as other groups have done. This strategy, implemented using a CCA with NMF approach, identified two meta-clusters as the model that best fit the data, and these two meta-clusters appear to have clinical relevance since cognitive decline is more rapid in MC-1. Importantly, we replicated this observation in an independent set of ROSMAP participants that were not used in the Discovery CCA analysis (**Fig. S2G** and **H**; **Table S1** and **S2**) and found consistent results in two other external sample collections (**Fig. S6**).

Our population structure captures an aspect of the molecular state of the brain that affects cognition in a manner that may be independent of the underlying pathologies that accumulate with older age (**Fig. 2**, **Table S2** and **S3**). While tau accumulation between the two groups is only marginally different, its biological impact as measured by the epigenomic perturbation score appears to be greater in MC-1 (**Table S4**): the impact of the same amount of tau pathology has a greater impact on the neuronal nucleus and, later, on the cognitive trajectory of MC-1 individuals.

Our epigenomic perturbation score is a summary measure of large-scale chromatin alterations(*19*). Although we cannot fully capture the state of chromatin with a single chromatin mark, our carefully selected mark does capture important changes in chromatin structure that relate to Tau pathology. Consistent with the results for quantitative measures of tau that are presented in **Table S4**, we see a skew towards higher Braak stages in MC-1. However, the population structure that we have found in sampling the three regions related to cognition that we selected to study is not driven by Braak stages. Further, the difference in cognitive decline between MC-1 and MC-2 remains significant after adjusting for Braak stage (𝑝𝑝_𝐹𝐹𝐹𝐹𝐹𝐹_ = 0.0001; **Table S3**).

As seen in **Fig. 2D and E**, large proportions of individuals from both MC-1 and MC-2 had AD at the time of death. Thus, we were not simply separating cognitively impaired and non-impaired individuals. Rather, we discriminate a subset of individuals that may be less resilient and more likely to decline for a given exposure: for example, our interaction analyses reveal that the *APOEε4* haplotype risk factor has a different effect size in MC-1 than in MC-2 brains. This elaborates the narrative that the effect of *APOEε4* can be influenced by other factors(*29*), including genetic ancestry, and that not all APOE genotypes are equivalent(*33*), although our analysis did not uncover significant differences in genotype frequencies between MC-1 And MC-2. Specifically, MC-1 subjects are impacted more strongly by this factor that contributes to the pathophysiological cascade of AD at different levels and leads to the accumulation of AD pathologies, then cognitive decline, the symptom of AD, and ultimately dementia. These results suggested that the MC-1/MC-2 partition captures some molecular process present in the transcriptome that globally renders MC-1 participants more likely to develop to cognitive decline; we therefore consider MC-1 individuals – who have a greater impact of risk factors (such as *APOEe4*) on AD risk and greater biological changes in relation to pathologic processes – as “vulnerable”. One interesting observation was that frontal white matter integrity (R2), which was previously associated with cognitive decline(*21*) was also more disrupted in MC-1 participants, suggesting that these gray-matter derived meta-clusters may be capturing a broader state of brain disruption that affects the aging brain.

The nature of the signature differentiating MC-1 and MC-2 is not well defined. It seems to contain genes that have previously been annotated as being members of gene signatures for a number of different neurodegenerative diseases. As such, the meta-clusters were not related to AD but may be an intrinsic property of certain brains. Consistent with this, there were large differences in cellular composition between MC-1 and MC-2, with fewer neurons and, interestingly, greater expression of the aged microglia gene set (HuMi-Aged) in MC-1. Since cognitively non-impaired individuals showed the same differences, these differences in cell type proportion and signatures were not occurring in the late phase of the disease, but we could not resolve whether they may have occurred during development or during the course of early brain aging.

Are there only two subtypes of aging brains? Given the heterogeneity in cognitive performance and wide variety of combinations of pathologic traits in ROSMAP subjects (1), we considered the possibility that we would uncover multiple subsets of subjects. However, differences in AD pathologic features were not the primary difference between our meta-clusters (**Fig. 2A** and **Table S2**), and two meta-clusters was the best empirical solution from the transcriptomic data. One limitation of the CCA approach is that it may de-prioritize data structure that are present in only one region as it seeks to achieve consensus over the three regions, minimizing region-specific effects. As shown in **Fig. S2A** and **G**, there is some difference in region-specific and meta-cluster assignments, so further work in individual regions is warranted to resolve local effects. The other two recent efforts to explore a subset of these data focused on individual regions and investigated a different question: how many subsets of brains existed among those that have a diagnosis of AD(*8, 9*). Thus, it did not consider the full spectrum of neocortical states in the older population (those participants with no or mild cognitive impairment that do not fulfill criteria for AD). While one of these efforts found no phenotypic differences among their clusters, the other found that inter-group differences driven by pathology, which is quite different from the difference in cognition-related changes that we have found with our meta-clustering approach. Overall, these approaches are likely to be complementary to the one that we deployed here, and further work can begin to integrate these different models. Another limitation of our analysis is that our large RNAseq dataset was generated over many years in different batches; this potential technical confounder was carefully considered and incorporated into our trait analyses to avoid over-correcting the data at earlier stages, such as when the expression residuals are calculated. Finally, an additional limitation of our study is that the ROSMAP participants are non-demented at the time of recruitment (usually in their mid-70’s) and had an average age of 89.6 years at the time of death. Thus, our results may not be generalizable to younger individuals.

Overall, we identified two major subsets of older brains that were related to biological processes influencing the impact of certain risk factors, such as *APOEε4*. These meta-clusters are meaningful because they capture ∼1% of the variance in aging-related cognitive decline, similar to neocortical lewy body burden which explains another 2.4% of variance, beyond what is explained by brain pathology and loss of white matter integrity (R2). Thus, if we understood the biological basis of this process, there may be an opportunity to mimic the effect that distinguishes the more resilient MC-2 individuals from the MC-1 individuals in which AD factors have a greater impact. Finding proxy markers for MC-1/MC-2 in peripheral blood or cerebrospinal fluid will be critical to enable to pursue the relevance of this classification in younger populations, in the course of AD and in clinical trials.

## MATERIALS AND METHODS

### Study Design

This study aimed to identify subsets of aging individuals using a data-driven approach and a large collection of individuals with RNA sequence data from three brain regions involved in cognitive circuits. Participants in this study were from the Religious Orders Study and Rush Memory and Aging Project (ROSMAP)(*4*), two clinical-pathologic cohort studies of aging and dementia, one from across the United States and the other from the greater Chicago area conducted by investigators at the Rush Alzheimer’s Disease Center. Transcriptome RNA sequencing data was accessed from postmortem brains including AC of 731 samples, DLPFC of 1,092 samples and PCC of 661 samples in ROSMAP. A total of 1,149 ROSMAP participants had transcriptomic data in at least one brain region. A subset of 459 participants had RNA-seq profiles in all three brain regions and served as the Discovery cohort (**Fig. 1A**). The remaining 690 participants had RNA-seq profiles in only one or two regions and were used as a Replication cohort. All participants are older and recruited free of known dementia (mean age at entry = 81.0 ± 7 (SD) years), are administered annual clinical and neuropsychological evaluations, and signed an Anatomical Gift Act allowing for brain autopsy at time of death. Written informed consent and a repository consent for data sharing were obtained from ROSMAP participants, and both studies were approved by an Institutional Review Board of Rush University Medical Center. ROSMAP data can be requested at https://www.radc.rush.edu.

### RNA sequencing data quantification, pre-processing and normalization

Data were generated in multiple batches from different sequencing centers and experimental protocols. RNA sequencing of dorsolateral prefrontal cortex was first done in 10 batches for 739 subjects at the Broad Institute. Samples were extracted using Qiagen’s miRNeasey Mini Kit (cat. no. 217004) and the RNase-Free DNase Set (cat. no. 79254). RNA was quantified using Nanodrop and RNA quality was evaluated using the BioAnalyzer (Agilent) as previously described(*34*). Samples with RNA Integrity Number (RIN) score greater than 5 and quantity threshold of 5μg were submitted for library construction. 569 subjects in 8 batches were processed using the dUTP method and 170 subjects in 2 batches were processed using the newer Illumina TruSeq method modified by The Broad Institute Genomics Platform. Sequencing was conducted using the Illumina HiSeq2000 with 101 bp paired end reads for a targeted coverage of 50 million paired reads. Subsequently, 124 subjects in 2 batches were sequenced at the New York Genome Center.

RNA-Seq library was prepared using the KAPA Stranded RNA-Seq Kit with RiboErase (Kapa Biosystems) according to the manufacturer’s instructions. Sequencing was carried out using the Illumina NovaSeq6000 sequence using 2×100bp cycles targeting 30 million reads per sample. Details of RNA sequencing was described previously(*34*). Moreover, 229 samples in a single batch were sequenced at the Rush Alzheimer’s Decease Center. RNA was extracted using Chemagic RNA tissue kit (Perkin Elmer, CMG-1212) and RNA Quality Number (RQN) was calculated using Fragment Analyzer (Agilent, DNF-471). 500ng total RNA was used to generate RNA-Seq library and rRNA was depleted with RiboGold (Illumina, 20020599). TruSeq stranded sequencing libraries (Illumina, 20020599) with custom unique dual indexes (IDT) were generated by the Zephyr G3 NGS workstation (Perkin Elmer). Libraries were normalized and sequenced on the Illumina NovaSeq 6000 with 2×150bp paired end at 40-50 million reads. Detailed RNA sequencing was described in Yu *et al.*, 2020(*34*). RNA sequencing of anterior caudate was done in 76 subjects in a single batch at the Broad Institute using the Illumina TruSeq method, as described above in DLPFC. Subsequently, 655 subjects in 2 batches were sequenced at the New York Genome Center using the same experimental protocols described in DLPFC. RNA sequencing of posterior cingulate cortex was done in 79 subjects in a single batch at the Broad Institute using the Illumina TruSeq method. Subsequently, 487 subjects in 2 batches were sequenced at the New York Genome Center and 95 subjects in a single batch were sequenced at the Rush Alzheimer’s Decease Center. The experimental protocols of each sequencing center are same as described in DLPFC.

Each brain region was pre-processed separately. We used gene-level transcription values derived from the RNA-seq data. The transcripts per million (TPM) values were used to measure the gene expression levels and subsequently normalized using Trimmed means of M-values (TMM) to create a frozen dataset that are available on Synapse (Synapse: syn25741873). Outlier samples were removed based on quantified expression profiles and lowly expressed genes with a median TPM less than 10 were filtered out to reduce the influence of technical noise.

To identify potentially important covariates that should be considered for adjustment for expression modeling, for example as in the DLPFC, we first looked at a distribution of participants by batch in **Fig. S7A** - which is generated from a MDS plot of gene transcriptions of TPM values - we observed that there was a difference between samples by batch. Next, when we explored correlation between various variables and first 20 principal components (PCs) from the gene transcript, we observed some of the technical covariates and pathologic variables were highly correlated with the first few PCs (**Fig. S7B**). We then used a forward selection method to identify covariates to be adjusted in the model. We started with a model including age, sex and batch and then successively added more technical variables if the technical variables are significantly associated with the PCs calculated from the residuals of the previous model. The selected variables for DLPFC were age at death, sex, and technical confounding factors which were batch, library size, percentage of coding bases, percentage of aligned reads, percentage of ribosomal bases, percentage of UTR base, median 5 prime to 3 prime bias, median CV coverage, pmi, and study index of ROS or MAP. Technical confounding factors for AC and PCC are very similar. To account for differences between samples, studies, experimental batch effects and unwanted RNA-sequencing specific technical variations, we adjusted for the selected variables in a linear regression and fitted for each transcript with the log2-transformed normalized TPM values as outcome. We used the residual from the transcript as our final transcription dataset. The residuals represent a quantitative trait capturing variability in transcripts outcome not captured by known demographics or technical factors. As a sanity check, when we explored the correlation between various variables and the first 20 PCs of residuals of the final model, the selected variables that we adjusted in the model were no longer associated with PCs of residuals (**Fig. S7C**). Adjusting for all these technical variables are more powerful than adjusting for batch alone. We adapted this approach from another reprocessing effort of the ROSMAP data(*35*).

We elected not to over-correct our data as adjusting for unknown surrogate variables calculated from the data could eliminate the biological effects that we are interested in (such as meta-clusters). This was a choice we made, and we mitigate the concern that population structure that we report could be spurious by (1) showing that this structure is related to meaningful clinical outcomes such as cognitive decline (which would be unlikely for a random technical factor), (2) we find a number of biological differences between the two meta-clusters that are not dependent on RNA (e.g. the epigenomic score and MRI measures, for example), and (3) we replicate our findings in an independent set of ROSMAP participants as well as other cohorts which underwent independent data pre-processing (**Fig. S6**). When we performed a surrogate variable analysis (SVA) using the R “sva” package(*36*) conditioned on age, sex, amyloid, tau, and batch effects to capture latent confounders for gene expression data, we did not find surrogate variables for the three regions.

### Statistical Analysis

We applied a sparse multiple canonical correlation analysis (CCA) in the Discovery cohort of individuals that have the residuals of normalized gene expression data in all three brain regions, in order to identify shared correlation structures across the three regions. CCA finds linear combinations of features that maximize the overall correlation across all regions. Formally, for 3 data sets, 𝑿𝑿 ∈ ℝ^𝑛𝑛×𝑝𝑝^, 𝒀𝒀 ∈ ℝ^𝑛𝑛×𝑞𝑞^, ***Z*** ∈ ℝ^𝑛𝑛×𝑟𝑟^, the multiple CCA finds projection vectors 𝒖𝒖, 𝒗𝒗, 𝒘𝒘 that maximize the sum of the pairwise correlations between 𝑗𝑗th canonical variates from the three brain regions 𝜌̂_𝑗𝑗_ = 𝑐𝑐𝑐𝑐𝑐𝑐̂𝑿𝑿𝒖𝒖_𝑗𝑗_, 𝒀𝒀𝒗𝒗_𝒋𝒋_ + 𝑐𝑐𝑐𝑐𝑐𝑐̂𝑿𝑿𝒖𝒖_𝑗𝑗_, 𝒁𝒁𝒘𝒘_𝒋𝒋_ + 𝑐𝑐𝑐𝑐𝑐𝑐̂𝒀𝒀𝒗𝒗_𝒋𝒋_, 𝒁𝒁𝒘𝒘_𝒋𝒋_. In other words, multiple CCA finds projection vectors 𝒖𝒖, 𝒗𝒗, 𝒘𝒘 that maximize max 𝒖𝒖^𝑻𝑻^𝑿𝑿^𝑻𝑻^𝒀𝒀𝒗𝒗 + 𝒖𝒖^𝑻𝑻^𝑿𝑿^𝑻𝑻^𝒁𝒁𝒘𝒘 + 𝒗𝒗^𝑻𝑻^𝒀𝒀^𝑻𝑻^𝒁𝒁𝒘𝒘 subject to 𝒖𝒖^𝑻𝑻^𝑿𝑿^𝑻𝑻^𝑿𝑿𝒖𝒖 ≤ 1, 𝒗𝒗^𝑻𝑻^𝒀𝒀^𝑻𝑻^𝒀𝒀𝒗𝒗 ≤ 1, 𝒘𝒘^𝑻𝑻^𝒁𝒁^𝑻𝑻^𝒁𝒁𝒘𝒘 ≤ 1. 𝑢𝑢,𝑣𝑣,𝑤𝑤

In our study, 𝑿𝑿, 𝒀𝒀, 𝒁𝒁 are the matrices of residual gene transcription of 𝑝𝑝, 𝑞𝑞, 𝑐𝑐 genes in DLPFC, AC, and PCC respectively from the same 𝑛𝑛 individuals, 𝒖𝒖_𝑗𝑗_, 𝒗𝒗_𝒋𝒋_, 𝒘𝒘_𝒋𝒋_ denote the 𝑗𝑗th column of penalized loadings or weights of DLPFC, AC, and PCC respectively that makes the sum to be maximized while satisfying constraints on 𝒖𝒖, 𝒗𝒗, 𝒘𝒘, and 𝑿𝑿𝒖𝒖, 𝒀𝒀𝒗𝒗, 𝒁𝒁𝒘𝒘 are called the canonical variates of DLPFC, AC, and PCC respectively which is a linear combination of the gene transcript using the given weights. Since 𝑝𝑝, 𝑞𝑞, 𝑐𝑐 were larger than 𝑛𝑛, we considered a sparse multiple CCA that finds linear combinations of features in 𝑿𝑿, 𝒀𝒀, 𝒁𝒁 that has large correlation but is also sparse in the variables used. One way to obtain penalized canonical variates is to assume 𝑿𝑿^𝑻𝑻^𝑿𝑿 = 𝒀𝒀^𝑻𝑻^𝒀𝒀 = 𝒁𝒁^𝑻𝑻^𝒁𝒁 =𝑰𝑰 as described in Witten and Tibshirani (2009)(*11*) and include penalties as follows:

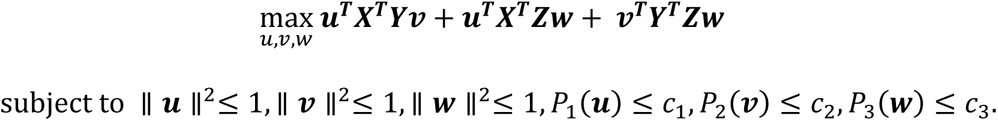

To determine the sparse parameters, we considered eleven sets of sparse parameters where sparse parameter for each brain region ranged from 2 to 20 increasing by 2 (i.e., (2,2,2), (4,4,4), …, (20,20,20): sparse parameters for DLPFC, AC, and PCC respectively) and used MultiCCA.permute function in R package PMA to select the sparse parameters. We considered the range of sparse parameters up to 20 in order to shrink the coefficient estimates towards zero and avoid selecting too many features. The optimal sparse parameters were (4,4,4) with a maximum sum of the pairwise correlations between first canonical variates from the three regions (𝜌̂_1_=1.14) (**Fig. S5A**). When we further explored the sum of the correlations for each canonical variates, ranging from 1^st^ to 6^th^ canonical variates, under various tuning parameter sets (**Fig. S5C** and **D**), sparse parameters set of (4,4,4) had overall large sum of the correlations. In **Fig. S5E**, we observed that around 30 genes were contributing to each canonical variate in three regions when we used a sparse CCA under the sparse parameters set of (4,4,4).

The number of canonical variates were selected based on the maximum eigenvalue ratio criterion(*37*). When we deployed sparse multiple CCA in the Discovery cohort of individuals, the sum of the pairwise correlations between each canonical variates from the three regions 𝜌̂_1_, …, 𝜌̂_𝑟𝑟𝑟𝑟𝑟𝑟𝑟𝑟_ were estimated (**Fig. S5A**). We considered a maximum of 𝑐𝑐𝑟𝑟𝑟𝑟𝑟𝑟 =10 canonical variates as in other high-dimensional data(*38*). We extract first 𝑘𝑘 canonical variates that maximize the ratio 𝜑𝜑̂_𝑗𝑗_ = 𝜌̂_𝑗𝑗_/𝜌̂_𝑗𝑗+1_ over the set of 𝑗𝑗 = 1, …, 𝑐𝑐𝑟𝑟𝑟𝑟𝑟𝑟 − 1. From the **Fig. S5B**, the ratio was maximized at 𝑗𝑗 =6 and we selected first six canonical variates in each region. We used the MultiCCA function in the R package PMA with sparse parameters set of (4,4,4) and standardized the data to have mean of zero and variance of one.

We subsequently deployed the k-means clustering to the first six canonical variates in each region returned from the sparse multiple CCA to identify subgroups of Discovery cohort of participants using the R package stats. To determine the optimal number of clusters in each region, we evaluated all possible models from 2 to 10 clusters, under three clustering methods (k-means clustering, hierarchical clustering and pam), and we evaluated the results using three internal clustering validation measures (connectivity measure, the Dunn index, and Silhouette width). As shown in **Fig. S3A**, for all three regions, two clusters of participants were optimal as the connectivity(*14*) was minimized. The Dunn index(*15*) and average silhouette width(*16*) were maximized under both k-means and hierarchical clustering. To validate the stability of our clusters, we used the clusterwise Jaccard bootstrap mean (CJBM)(*17*) and considered two to five clusters in each region under the k-means and hierarchical clustering methods. As shown in **Fig. S3B**, 2 clusters obtained through the k-means clustering were the most stable across clusters in all three regions because both clusters had CJBM above 0.85. Therefore, we have prioritized two clusters of participants defined by using k-means clustering as our optimal model.

The two clusters of Discovery participants identified from different brain regions were then integrated across the brain regions into two meta-clusters through a non-negative matrix factorization (NMF)(*12*) approach. Specifically, the cluster assignments of each subject in Discovery cohort from three different regions were first combined into a matrix. The NMF model was then fitted on the matrix for each subject and factorized the matrix into a basis matrix representing the projection of original clusters from each region to the meta-clusters and a coefficient matrix representing the projection of the individual participants to meta-clusters, showing the meta-clusters membership of the individual participants. The number of meta-clusters 𝑘𝑘 that is too low could merge unrelated clusters, while 𝑘𝑘 that is too high could potentially fail to integrate the related clusters. In the ideal case, each region-specific cluster will perfectly project onto one of the meta-clusters. For Discovery cohort of individuals, the optimal number of meta-clusters was two clusters of participants based on the Cophenetic coefficient(*39*) which represents the global clustering robustness score, where higher value indicates more robust partition (**Fig. S2C**). The stability of the clusters was assessed through the consensus matrix and two meta-clusters were stable as the subdivision was evident (**Fig. S2D**). The NMF results for two meta-clusters are shown in **Fig. S2A.**

To partition the remaining Replication cohort of individuals that have data in only one or two of the three regions, the same canonical weights 𝒖𝒖, 𝒗𝒗, 𝒘𝒘 obtained from sparse multiple CCA in the Discovery cohort were given to the features in the Replication cohort, returning canonical variates for each region. The subgroups of participants were then identified by assigning their canonical variates to the nearest cluster centroid found in the Discovery cohort, leading to two clusters of participants for each region (**Fig. S1**). Through NMF, the resulting clusters were integrated into two stable meta-clusters of participants in the Replication cohort (**Fig. S2G)**.

To assess whether clinico-pathologic traits were different between two transcriptomically-defined meta-clusters, a regression model was fitted on Discovery and Replication cohorts of individuals separately and evaluated the effect of meta-clusters on the trait. A linear regression model was fitted on a continuous trait and a logistic regression model was fitted on a binary trait after adjusting for age and sex. We analyzed the square root of β-amyloid and tau to transform it to follow normal distributions for modeling. We then conducted a meta-analysis of the two association studies obtained from Discovery and Replication cohorts through fixed-effects model with inverse-variance weighted average method. We tested the effect of meta-clusters in order to increase the statistical power and created **Fig. 2A**, **Fig. S6A, Table S2-S6 and S10**.

To further investigate whether the association between two clinico-pathologic traits vary by two meta-clusters, two separate regression models were fitted on individuals within each meta-cluster and evaluated the association between two traits, for each Discovery and Replication cohorts separately. A linear regression model was fitted on a continuous trait and a logistic regression model was fitted on a binary trait after adjusting for age and sex. We then conducted a meta-analysis of the two associations studies from Discovery and Replication cohorts for each meta-clusters separately. We tested the difference in the association between two meta-clusters by fitting a single fixed-effects meta-regression model with meta-clusters as a moderator using the R package metafor and created **Fig. S6B**, **Table S7**, **S8** and **S11**.

While we assessed the difference in clinico-pathologic traits between MC-1 and MC-2 using the meta-analysis of association results from Discovery and Replication cohorts, for visualization purposes only, we simply evaluated the difference for 1,149 ROSMAP participants with transcriptomic data in one or more brain regions to compute the p-values in **Fig. 2B**, **D** and **E, Table 2** and **Fig. S4**. The mean difference in a continuous trait between MC-1 and MC-2 were assessed by using an independent t-test or the non-parametric equivalent Wilcoxon rank sum test. Proportions of measured variables by two meta-clusters were assessed by chi-square tests. The normality of the data distribution was determined using the Shapiro-Wilk test.

To estimate age at AD onset in MC-1 and MC-2, we used the extended accelerated failure time model(*40*) as described in Wilson *et al.*, 2021(*41*). ROSMAP participants were all non-demented at study entry which can bias estimates of survival parameters due left-truncation. To adjust for the left-truncation, we used the extended accelerated failure time model with generalized gamma distribution for a baseline age distribution(*41*) controlling for sex and education using the flexsurvreg function in R package flexsurv(*40*) and estimated age at AD onset and cumulative risk of AD onset for MC-1 and MC-2 as presented in **Fig. 7** and **Table S9**, respectively.

Of note, all statistical analyses were performed using R 4.1.3 (https://www.r-project.org/). Benjamini & Hochberg’s false discovery rate (FDR) of 0.05 was used as a threshold for statistical significance.

## Supplementary Materials

### Materials and Methods

Fig. S1. K-means clustering results for Discovery and Replication cohorts in each brain region.

Fig. S2. NMF-based clustering on obtaining meta-cluster assignment from clusters of participants identified from each region and consensus analysis of the Discovery and Replication cohorts.

Fig. S3. Performance of clustering methods under various number of clusters for Discovery cohort within each brain region.

Fig. S4. Box plot of MRI-derived frontal white matter R2 by meta-cluster assignment.

Fig. S5. Sum of the pairwise correlations from the three brain regions under various sparse parameters by canonical variates.

Fig. S6. Validation study in Mount Sinai Brain Bank and Mayo Clinic. Fig. S7. Exploring covariates to be adjusted for expression modeling.

Table S1. Distribution of participants by clusters obtained from each brain region and meta-clusters.

Table S2. Association of traits with meta-cluster assignment, after adjusting for age and sex.

Table S3. Meta-analysis association of cognitive decline between MC-1 and MC-2, after adjusting for covariate(s) in addition to age and sex.

Table S4. Meta-analysis association of other factors between MC-1 and MC-2, after adjusting for age and sex.

Table S5. Meta-analysis of association of factor between MC-1 and MC-2 among participants with pathological AD, among those without pathological AD, among those with AD dementia, or among those without AD dementia, after adjusting for age and sex.

Table S6. Meta-analysis association of AD-related gene between MC-1 and MC-2, after adjusting for age and sex.

Table S7. Strength of evidence for interaction of meta-cluster assignment with predictor on outcome, after adjusting for age and sex.

Table S8. Strength of evidence for interaction of meta-cluster assignment with APOEε4 on cognitive decline among participants with pathological AD, among those without pathological AD, among those with AD dementia, or among those without AD dementia, after adjusting for age and sex.

Table S9. Estimated cumulative risk of AD dementia onset in MC-1 and MC-2, adjusting for sex and education in ROSMAP (× 10−2).

Table S10. Meta-analysis association of pathologic diagnosis of AD between MC-1 and MC-2 in ROSMAP, MSBB and Mayo, after adjusting for age and sex.

Table S11. Strength of evidence for interaction of meta-cluster assignment with APOEε4 on pathologic diagnosis of AD, in ROSMAP, MSBB and Mayo, after adjusting for age and sex.

## Acknowledgements

**Acknowledgements:** We would like to thank the participants and working group of the Religious Order Study, the Memory and Aging Project. Funding: This study has been supported by the National Institutes of Health grants P30AG10161 (D.A.B), R01AG15819 (D.A.B), R01AG17917 (D.A.B), U01AG61356 (D.A.B), R01AG036836 (P.L.D.), U01AG046152 (P.L.D. and D.A.B), and TAME-AD (The Thompson Family Foundation Program for Accelerated Medicine Exploration in Alzheimer’s Disease and Related Disorders of the Nervous System). **Author contributions**: A.J.L., H.K, and P.L.D. created the study concept, designed the study, and interpreted the experiments. A.J.L. and P.L.D. drafted the manuscript. A.J.L. performed and analyzed the experiments. H.K performed experiments in Fig. 2C. Y.M analyzed the monocyte RNA-seq data analyses. All authors contributed to data acquisition and interpretation and critically revised the manuscript. **Competing interests**: The authors declare that they have no competing interests. **Data and materials availability**: The normalized RNA-seq data from the three brain regions are available via the AD Knowledge Portal (https://adknowledgeportal.org). The AD Knowledge Portal is a platform for accessing data, analyses, and tools generated by the Accelerating Medicines Partnership (AMP-AD) Target Discovery Program and other National Institute on Aging (NIA)-supported programs to enable open-science practices and accelerate translational learning. The data, analyses and tools are shared early in the research cycle without a publication embargo on secondary use. Data is available for general research use according to the following requirements for data access and data attribution (https://adknowledgeportal.org/DataAccess/Instructions). Content described in this manuscript are available on Synapse (Synapse: syn25741873).

