## Supplementary Materials for "Multi-region brain transcriptomes uncover two subtypes of aging individuals with differences in Alzheimer risk and the impact of *APOEε4*"

**One Sentence Summary:** There are two types of aging brains, with one being more vulnerable to *APOEε4* and subsequent neuronal dysfunction and cognitive loss.

**Abstract**: The heterogeneity of the older population suggests the existence of subsets of individuals which share certain brain molecular features and respond differently to risk factors for Alzheimer's disease, but this population structure remains poorly defined. Here, we performed an unsupervised clustering of individuals with multi-region brain transcriptomes to assess whether a broader approach, simultaneously considering data from multiple regions involved in cognition would uncover such subsets. We implemented a canonical correlation-based analysis in a Discovery cohort of 459 participants from two longitudinal studies of cognitive aging that have RNA sequence profiles in three brain regions. 690 additional participants that have data in only one or two of these regions were used in the Replication effort. These clustering analyses identified two meta-clusters, MC-1 and MC-2. The two sets of participants differ primarily in their trajectories of cognitive decline, with MC-2 having a delay of 3 years to the median age of incident dementia. This is due, in part, to a greater impact of tau pathology on neuronal chromatin architecture and to broader brain changes including greater loss of white matter integrity in MC-1. Further evidence of biological differences includes a significantly larger impact of *APOEε4* risk on cognitive decline in MC-1. These findings suggest that our proposed population structure captures an aspect of the more distributed molecular state of the aging brain that either enhances the effect of risk factors in MC-1 or of protective effects in MC-2. These observations may inform the design of therapeutic development efforts and of trials as both become increasingly more targeted molecularly.

**Supplementary Materials**

Materials and Methods

Fig. S1. K-means clustering results for Discovery and Replication cohorts in each brain region.

Fig. S2. NMF-based clustering on obtaining meta-cluster assignment from clusters of participants identified from each region and consensus analysis of the Discovery and Replication cohorts.

Fig. S3. Performance of clustering methods under various number of clusters for Discovery cohort within each brain region.

Fig. S4. Box plot of MRI-derived frontal white matter $R_2$ by meta-cluster assignment.

Fig. S5. Sum of the pairwise correlations from the three brain regions under various sparse parameters by canonical variates.

Fig. S6. Validation study in Mount Sinai Brain Bank and Mayo Clinic.

Fig. S7. Exploring covariates to be adjusted for expression modeling.

Table S1. Distribution of participants by clusters obtained from each brain region and meta-clusters.

Table S2. Association of traits with meta-cluster assignment, after adjusting for age and sex.

Table S3. Meta-analysis association of cognitive decline between MC-1 and MC-2, after adjusting for covariate(s) in addition to age and sex.

Table S4. Meta-analysis association of other factors between MC-1 and MC-2, after adjusting for age and sex.

Table S5. Meta-analysis of association of factor between MC-1 and MC-2 among participants with pathological AD, among those without pathological AD, among those with AD dementia, or among those without AD dementia, after adjusting for age and sex.

Table S6. Meta-analysis association of AD-related gene between MC-1 and MC-2, after adjusting for age and sex.

Table S7. Strength of evidence for interaction of meta-cluster assignment with predictor on outcome, after adjusting for age and sex.

Table S8. Strength of evidence for interaction of meta-cluster assignment with *APOEε4* on cognitive decline among participants with pathological AD, among those without pathological AD, among those with AD dementia, or among those without AD dementia, after adjusting for age and sex.

Table S9. Estimated cumulative risk of AD dementia onset in MC-1 and MC-2, adjusting for sex and education in ROSMAP ($\times 10^{-2}$).

Table S10. Meta-analysis association of pathologic diagnosis of AD between MC-1 and MC-2 in ROSMAP, MSBB and Mayo, after adjusting for age and sex.

Table S11. Strength of evidence for interaction of meta-cluster assignment with *APOEε4* on pathologic diagnosis of AD, in ROSMAP, MSBB and Mayo, after adjusting for age and sex.

Giedraitis, A. Kawalia, S. Li, R. M. Huebinger, L. Kilander, S. Moebus, I. Hernandez, M. I. Kamboh, R. Brundin, J. Turton, Q. Yang, M. J. Katz, L. Concari, J. Lord, A. S. Beiser, C. D. Keene, S. Helisalmi, I. Kloszewska, W. A. Kukull, A. M. Koivisto, A. Lynch, L. Tarraga, E. B. Larson, A. Haapasalo, B. Lawlor, T. H. Mosley, R. B. Lipton, V. Solfrizzi, M. Gill, W. T. Longstreth, Jr., T. J. Montine, V. Frisardi, M. Diez-Fairen, F. Rivadeneira, R. C. Petersen, V. Deramecourt, I. Alvarez, F. Salani, A. Ciaramella, E. Boerwinkle, E. M. Reiman, N. Fievet, J. I. Rotter, J. S. Reisch, O. Hanon, C. Cupidi, A. G. Andre Uitterlinden, D. R. Royall, C. Dufouil, R. G. Maletta, I. de Rojas, M. Sano, A. Brice, R. Cecchetti, P. S. George-Hyslop, K. Ritchie, M. Tsolaki, D. W. Tsuang, B. Dubois, D. Craig, C. K. Wu, H. Soininen, D. Avramidou, R. L. Albin, L. Fratiglioni, A. Germanou, L. G. Apostolova, L. Keller, M. Koutroumani, S. E. Arnold, F. Panza, O. Gkatzima, S. Asthana, D. Hannequin, P. Whitehead, C. S. Atwood, P. Caffarra, H. Hampel, I. Quintela, A. Carracedo, L. Lannfelt, D. C. Rubinsztein, L. L. Barnes, F. Pasquier, L. Frolich, S. Barral, B. McGuinness, T. G. Beach, J. A. Johnston, J. T. Becker, P. Passmore, E. H. Bigio, J. M. Schott, T. D. Bird, J. D. Warren, B. F. Boeve, M. K. Lupton, J. D. Bowen, P. Proitsi, A. Boxer, J. F. Powell, J. R. Burke, J. S. K. Kauwe, J. M. Burns, M. Mancuso, J. D. Buxbaum, U. Bonuccelli, N. J. Cairns, A. McQuillin, C. Cao, G. Livingston, C. S. Carlson, N. J. Bass, C. M. Carlsson, J. Hardy, R. M. Carney, J. Bras, M. M. Carrasquillo, R. Guerreiro, M. Allen, H. C. Chui, E. Fisher, C. Masullo, E. A. Crocco, C. DeCarli, G. Bisceglio, M. Dick, L. Ma, R. Duara, N. R. Graff-Radford, D. A. Evans, A. Hodges, K. M. Faber, M. Scherer, K. B. Fallon, M. Riemenschneider, D. W. Fardo, R. Heun, M. R. Farlow, H. Kolsch, S. Ferris, M. Leber, T. M. Foroud, I. Heuser, D. R. Galasko, I. Giegling, M. Gearing, M. Hull, D. H. Geschwind, J. R. Gilbert, J. Morris, R. C. Green, K. Mayo, J. H. Growdon, T. Feulner, R. L. Hamilton, L. E. Harrell, D. Drichel, L. S. Honig, T. D. Cushion, M. J. Huentelman, P. Hollingworth, C. M. Hulette, B. T. Hyman, R. Marshall, G. P. Jarvik, A. Meggy, E. Abner, G. E. Menzies, L. W. Jin, G. Leonenko, L. M. Real, G. R. Jun, C. T. Baldwin, D. Grozeva, A. Karydas, G. Russo, J. A. Kaye, R. Kim, F. Jessen, N. W. Kowall, B. Vellas, J. H. Kramer, E. Vardy, F. M. LaFerla, K. H. Jockel, J. J. Lah, M. Dichgans, J. B. Leverenz, D. Mann, A. I. Levey, S. Pickering-Brown, A. P. Lieberman, N. Klopp, K. L. Lunetta, H. E. Wichmann, C. G. Lyketsos, K. Morgan, D. C. Marson, K. Brown, F. Martiniuk, C. Medway, D. C. Mash, M. M. Nothen, E. Masliah, N. M. Hooper, W. C. McCormick, A. Daniele, S. M. McCurry, A. Bayer, A. N. McDavid, J. Gallacher, A. C. McKee, H. van den Bussche, M. Mesulam, C. Brayne, B. L. Miller, S. Riedel-Heller, C. A. Miller, J. W. Miller, A. Al-Chalabi, J. C. Morris, C. E. Shaw, A. J. Myers, J. Wiltfang, S. O'Bryant, J. M. Olichney, V. Alvarez, J. E. Parisi, A. B. Singleton, H. L. Paulson, J. Collinge, W. R. Perry, S. Mead, E. Peskind, D. H. Cribbs, M. Rossor, A. Pierce, N. S. Ryan, W. W. Poon, B. Nacmias, H. Potter, S. Sorbi, J. F. Quinn, E. Sacchinelli, A. Raj, G. Spalletta, M. Raskind, C. Caltagirone, P. Bossu, M. D. Orfei, B. Reisberg, R. Clarke, C. Reitz, A. D. Smith, J. M. Ringman, D. Warden, E. D. Roberson, G. Wilcock, E. Rogaeva, A. C. Bruni, H. J. Rosen, M. Gallo, R. N. Rosenberg, Y. Ben-Shlomo, M. A. Sager, P. Mecocci, A. J. Saykin, P. Pastor, M. L. Cuccaro, J. M. Vance, J. A. Schneider, L. S. Schneider, S. Slifer, W. W. Seeley, A. G. Smith, J. A. Sonnen, S. Spina, R. A. Stern, R. H. Swerdlow, M. Tang, R. E. Tanzi, J. Q. Trojanowski, J. C. Troncoso, V. M. Van Deerlin, L. J. Van Eldik, H. V. Vinters, J. P. Vonsattel, S. Weintraub, K. A. Welsh-Bohmer, K. C. Wilhelmsen, J. Williamson, T. S. Wingo, R. L. Woltjer, C. B. Wright, C. E. Yu, L. Yu, Y. Saba, A. Pilotto, M. J. Bullido, O. Peters, P. K. Crane, D. Bennett, P. Bosco, E. Coto, V. Boccardi, P. L. De Jager, A. Lleo, N. Warner, O. L. Lopez, M. Ingelsson, P. Deloukas, C. Cruchaga, C. Graff, R. Gwilliam, M. Fornage, A. M. Goate, P. Sanchez-Juan, P. G. Kehoe, N. Amin, N. Ertekin-Taner, C. Berr, S. Debette, S. Love, L. J. Launer, S. G. Younkin, J. F. Dartigues, C. Corcoran, M. A. Ikram, D. W. Dickson, G. Nicolas, D. Campion, J. Tschanz, H. Schmidt, H. Hakonarson, J. Clarimon, R. Munger, R. Schmidt, L. A. Farrer, C. Van Broeckhoven, C. O. D. M, A. L. DeStefano, L. Jones, J. L. Haines, J. F. Deleuze, M. J. Owen, V. Gudnason, R. Mayeux, V. Escott-Price, B. M. Psaty, A. Ramirez, L. S. Wang, A. Ruiz, C. M. van Duijn, P. A. Holmans, S. Seshadri, J. Williams, P. Amouyel, G. D. Schellenberg, J. C. Lambert, M. A. Pericak-Vance, C. Alzheimer Disease Genetics, I. European Alzheimer's Disease, H. Cohorts for, C. Aging Research in Genomic Epidemiology, Genetic, P. Environmental Risk in Ad/Defining Genetic, C. Environmental Risk for Alzheimer's Disease, Genetic meta-analysis of diagnosed Alzheimer's disease identifies new risk loci and implicates Abeta, tau, immunity and lipid processing. *Nat Genet* **51**, 414-430 (2019).

24. J. A. Mortimer, D. A. Snowdon, W. R. Markesbery, The effect of APOE-epsilon4 on dementia is mediated by Alzheimer neuropathology. *Alzheimer Dis Assoc Disord* **23**, 152-157 (2009).
25. M. Kanehisa, S. Goto, Y. Sato, M. Furumichi, M. Tanabe, KEGG for integration and interpretation of large-scale molecular data sets. *Nucleic Acids Res* **40**, D109-114 (2012).

26. Y. Zhong, Y. W. Wan, K. Pang, L. M. Chow, Z. Liu, Digital sorting of complex tissues for cell type-specific gene expression profiles. *BMC Bioinformatics* **14**, 89 (2013).
27. X. Wang, M. Allen, S. Li, Z. S. Quicksall, T. A. Patel, T. P. Carnwath, J. S. Reddy, M. M. Carrasquillo, S. J. Lincoln, T. T. Nguyen, K. G. Malphrus, D. W. Dickson, J. E. Crook, Y. W. Asmann, N. Ertekin-Taner, Deciphering cellular transcriptional alterations in Alzheimer's disease brains. *Mol Neurodegener* **15**, 38 (2020).
28. M. Olah, E. Patrick, A. C. Villani, J. Xu, C. C. White, K. J. Ryan, P. Piehowski, A. Kapasi, P. Nejad, M. Cimpean, S. Connor, C. J. Yung, M. Frangieh, A. McHenry, W. Elyaman, V. Petyuk, J. A. Schneider, D. A. Bennett, P. L. De Jager, E. M. Bradshaw, A transcriptomic atlas of aged human microglia. *Nat Commun* **9**, 539 (2018).
29. Y. Ma, L. Yu, M. Olah, R. Smith, S. R. Oatman, M. Allen, E. Pishva, B. Zhang, V. Menon, N. Ertekin-Taner, K. Lunnon, D. A. Bennett, H. U. Klein, P. L. De Jager, Epigenomic features related to microglia are associated with attenuated effect of APOE epsilon4 on Alzheimer's disease risk in humans. *Alzheimers Dement* **18**, 688-699 (2022).
30. M. Wang, N. D. Beckmann, P. Roussos, E. Wang, X. Zhou, Q. Wang, C. Ming, R. Neff, W. Ma, J. F. Fullard, M. E. Hauberg, J. Bendl, M. A. Peters, B. Logsdon, P. Wang, M. Mahajan, L. M. Mangravite, E. B. Dammer, D. M. Duong, J. J. Lah, N. T. Seyfried, A. I. Levey, J. D. Buxbaum, M. Ehrlich, S. Gandy, P. Katsel, V. Haroutunian, E. Schadt, B. Zhang, The Mount Sinai cohort of large-scale genomic, transcriptomic and proteomic data in Alzheimer's disease. *Sci Data* **5**, 180185 (2018).
31. M. Allen, M. M. Carrasquillo, C. Funk, B. D. Heavner, F. Zou, C. S. Younkin, J. D. Burgess, H. S. Chai, J. Crook, J. A. Eddy, H. Li, B. Logsdon, M. A. Peters, K. K. Dang, X. Wang, D. Serie, C. Wang, T. Nguyen, S. Lincoln, K. Malphrus, G. Bisceglio, M. Li, T. E. Golde, L. M. Mangravite, Y. Asmann, N. D. Price, R. C. Petersen, N. R. Graff-Radford, D. W. Dickson, S. G. Younkin, N. Ertekin-Taner, Human whole genome genotype and transcriptome data for Alzheimer's and other neurodegenerative diseases. *Sci Data* **3**, 160089 (2016).
32. M. Wang, P. Roussos, A. McKenzie, X. Zhou, Y. Kajiwara, K. J. Brennand, G. C. De Luca, J. F. Crary, P. Casaccia, J. D. Buxbaum, M. Ehrlich, S. Gandy, A. Goate, P. Katsel, E. Schadt, V. Haroutunian, B. Zhang, Integrative network analysis of nineteen brain regions identifies molecular signatures and networks underlying selective regional vulnerability to Alzheimer's disease. *Genome Med* **8**, 104 (2016).
33. E. M. Reiman, J. F. Arboleda-Velasquez, Y. T. Quiroz, M. J. Huentelman, T. G. Beach, R. J. Caselli, Y. Chen, Y. Su, A. J. Myers, J. Hardy, J. Paul Vonsattel, S. G. Younkin, D. A. Bennett, P. L. De Jager, E. B. Larson, P. K. Crane, C. D. Keene, M. I. Kamboh, J. K. Kofler, L. Duque, J. R. Gilbert, H. E. Gwirtsman, J. D. Buxbaum, D. W. Dickson, M. P. Frosch, B. F. Ghetti, K. L. Lunetta, L. S. Wang, B. T. Hyman, W. A. Kukull, T. Foroud, J. L. Haines, R. P. Mayeux, M. A. Pericak-Vance, J. A. Schneider, J. Q. Trojanowski, L. A. Farrer, G. D. Schellenberg, G. W. Beecham, T. J. Montine, G. R. Jun, C. Alzheimer's Disease Genetics, Exceptionally low likelihood of Alzheimer's dementia in APOE2 homozygotes from a 5,000-person neuropathological study. *Nat Commun* **11**, 667 (2020).
34. L. Yu, S. Tasaki, J. A. Schneider, K. Arfanakis, D. M. Duong, A. P. Wingo, T. S. Wingo, N. Kearns, G. R. J. Thatcher, N. T. Seyfried, A. I. Levey, P. L. De Jager, D. A. Bennett, Cortical Proteins Associated With Cognitive Resilience in Community-Dwelling Older Persons. *JAMA Psychiatry* **77**, 1172-1180 (2020).
35. Y. W. Wan, R. Al-Ouran, C. G. Mangleburg, T. M. Perumal, T. V. Lee, K. Allison, V. Swarup, C. C. Funk, C. Gaiteri, M. Allen, M. Wang, S. M. Neuner, C. C. Kaczorowski, V. M. Philip, G. R. Howell, H. Martini-Stoica, H. Zheng, H. Mei, X. Zhong, J. W. Kim, V. L. Dawson, T. M. Dawson, P. C. Pao, L. H. Tsai, J. V. Haure-Mirande, M. E. Ehrlich, P. Chakrabarty, Y. Levites, X. Wang, E. B. Dammer, G. Srivastava, S. Mukherjee, S. K. Sieberts, L. Omberg, K. D. Dang, J. A. Eddy, P. Snyder, Y. Chae, S. Amberkar, W. Wei, W. Hide, C. Preuss, A. Ergun, P. J. Ebert, D. C. Airey, S. Mostafavi, L. Yu, H. U. Klein, C. Accelerating Medicines Partnership-Alzheimer's Disease, G. W. Carter, D. A. Collier, T. E. Golde, A. I. Levey, D. A. Bennett, K. Estrada, T. M. Townsend, B. Zhang, E. Schadt, P. L. De Jager, N. D. Price, N. Ertekin-Taner, Z. Liu, J. M. Shulman, L. M. Mangravite, B. A. Logsdon, Meta-Analysis of the Alzheimer's Disease Human Brain Transcriptome and Functional Dissection in Mouse Models. *Cell Rep* **32**, 107908 (2020).
36. J. T. Leek, J. D. Storey, Capturing heterogeneity in gene expression studies by surrogate variable analysis. *PLoS Genet* **3**, 1724-1735 (2007).
37. B. An, Guo, J., Wang, H., Multivariate regression shrinkage and selection by canonical correlation analysis. *Computational Statistics & Data Analysis* **62**, 93-107 (2013).
38. I. Wilms, C. Croux, Robust sparse canonical correlation analysis. *BMC Syst Biol* **10**, 72 (2016).

39. J. P. Brunet, P. Tamayo, T. R. Golub, J. P. Mesirov, Metagenes and molecular pattern discovery using matrix factorization. *Proc Natl Acad Sci U S A* **101**, 4164-4169 (2004).
40. C. H. Jackson, flexsurv: A Platform for Parametric Survival Modeling in R. *J Stat Softw* **70**, (2016).
41. R. S. Wilson, T. Wang, L. Yu, F. Grodstein, D. A. Bennett, P. A. Boyle, Cognitive Activity and Onset Age of Incident Alzheimer Disease Dementia. *Neurology* **97**, e922-e929 (2021).

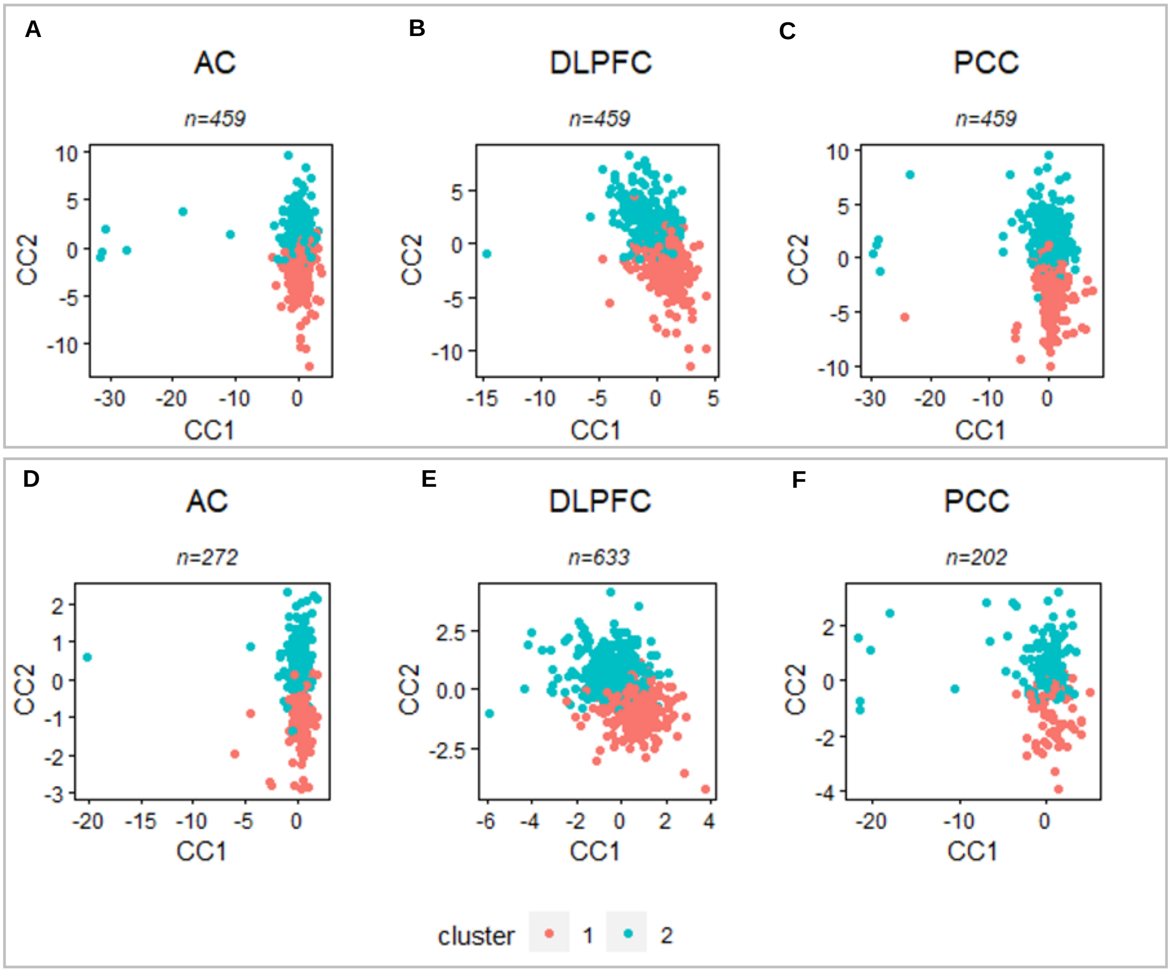


**Fig. S1. K-means clustering results for Discovery and Replication cohorts in each brain region**. Scatter plots of the first and second canonical variates returned from deploying sparse multiple CCA show two subgroups of (**A-C**) Discovery and (**D-F**) Replication participants in AC, DLPFC, and PCC respectively identified from applying k-means clustering. *Discovery* participants with an RNA-seq profile in all 3 regions; *Replication* those participants with an RNA-seq profile in only one or two regions; *AC* anterior caudate; *DLPFC* dorsolateral prefrontal cortex; *PCC* posterior cingulate cortex.

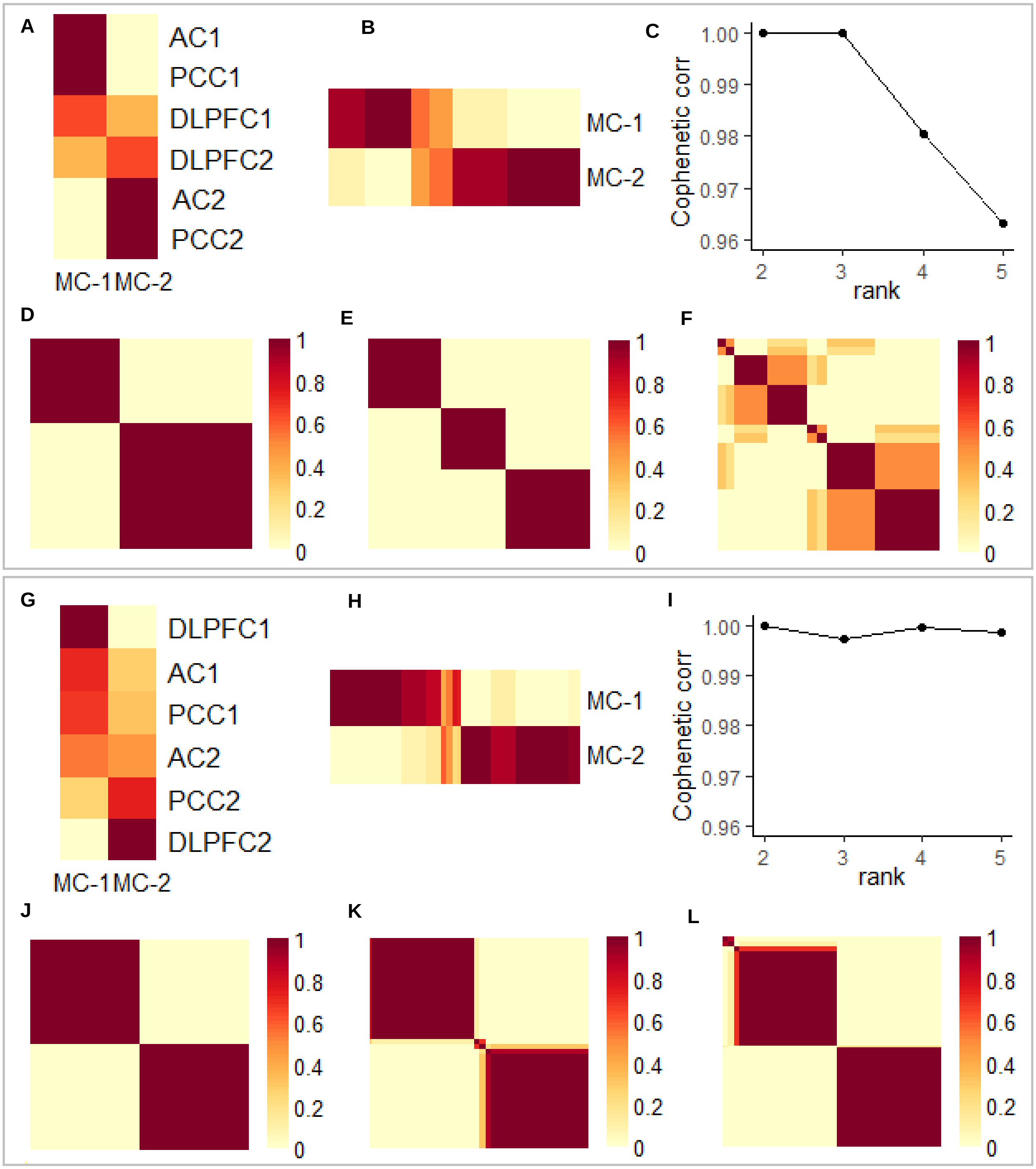


**Fig. S2. NMF-based clustering on obtaining meta-cluster assignment from clusters of participants identified from each region and consensus analysis of the Discovery and Replication cohorts.** (**A, G**) Basis matrices represent the projection of clusters obtained from each region in isolation (y-axis) to the meta-clusters (x-axis) in Discovery and Replication cohorts, respectively. (**B, H**) Coefficient matrix represents the projection of the Discovery or Replication participants (x-axis) respectively to meta-clusters (y-axis). (**C, I**) Cophenetic correlation coefficients (y-axis) over a range of two to five clusters (x-axis) in the Discovery and Replication cohorts, respectively. The cophenetic correlation coefficient indicates the robustness of the clusters, where higher value indicates more robust partition. (**D-F, J-L**) Consensus matrices for two to five clusters in the Discovery and Replication cohorts, respectively. The consensus matrix indicates the stability of the clusters, where evident subdivision indicates more stable clusters. The NMF consensus analysis revealed two meta-clusters were robust and stable clustering for both Discovery and Replication cohorts. *Discovery* participants with an RNA-seq profile in all 3 regions; *Replication* participants with an RNA-seq profile in only one or two regions; *MC* meta-cluster; *AC* anterior caudate; *DLPFC* dorsolateral prefrontal cortex; *PCC* posterior cingulate cortex.

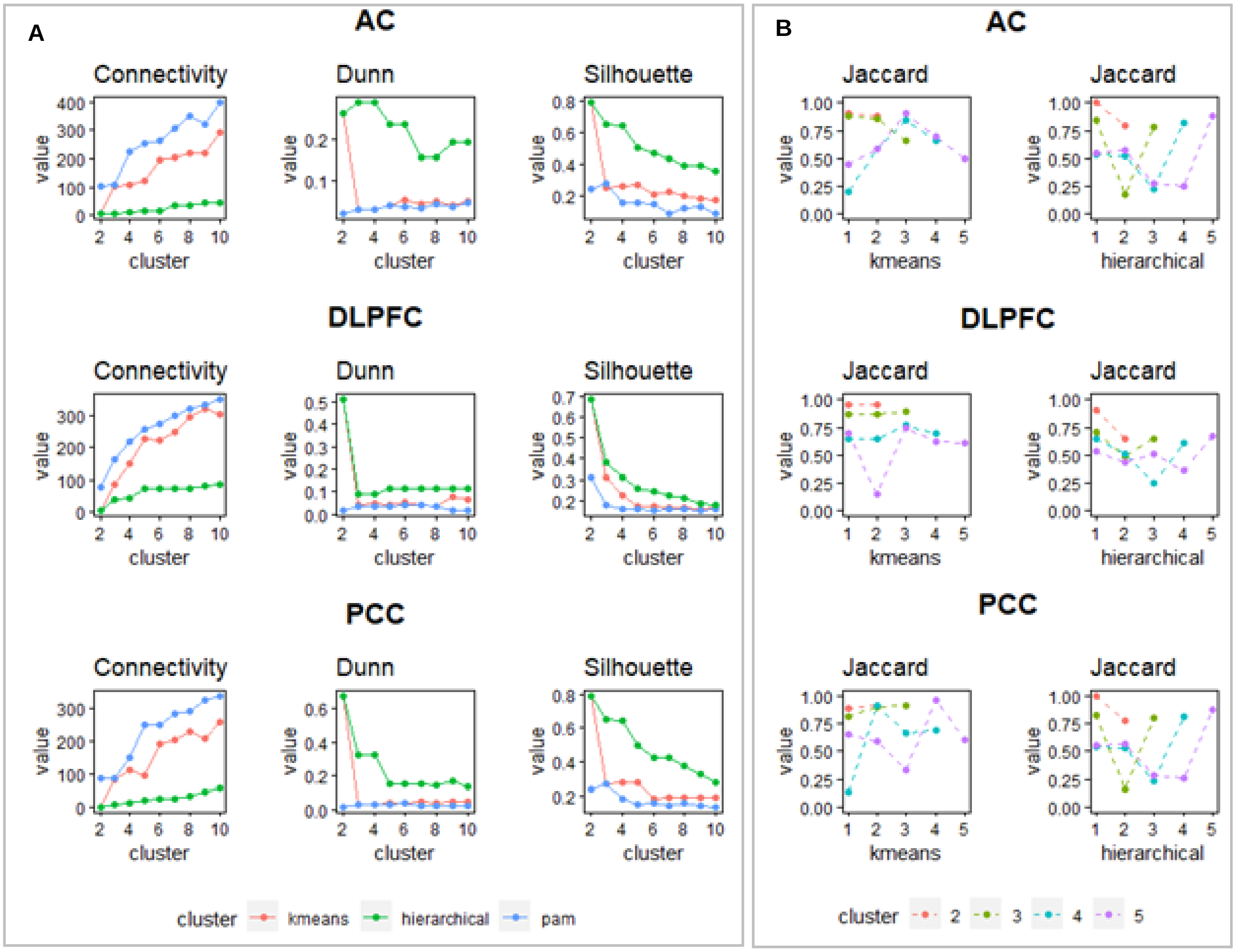


**Fig. S3. Performance of clustering methods under various number of clusters for Discovery cohort within each brain region.** (**A**) Three clustering methods (k-means clustering, hierarchical clustering, and partitioning around medoids (PAM)) with a range of models from 2 to 10 clusters were compared in each region. Three internal clustering validation measures (connectivity measure, the Dunn index, and Silhouette width) were used to evaluate the results. For all three regions, two clusters of participants were optimal as the connectivity was minimized and the Dunn index with average silhouette width were maximized under both k-means and hierarchical clustering. (**B**) For the clusters obtained through the k-means and hierarchical clustering, clustering stability was evaluated through clusterwise Jaccard bootstrap mean (CJBM) which measures the Jaccard similarities of the original clusters to the most similar clusters in the 1,000 bootstrapped samples with a range of models from 2 to 5 clusters in each region. For all three brain regions, two clusters of participants obtained through the k-means clustering were the most stable with both clusters having CJBM above 0.85.

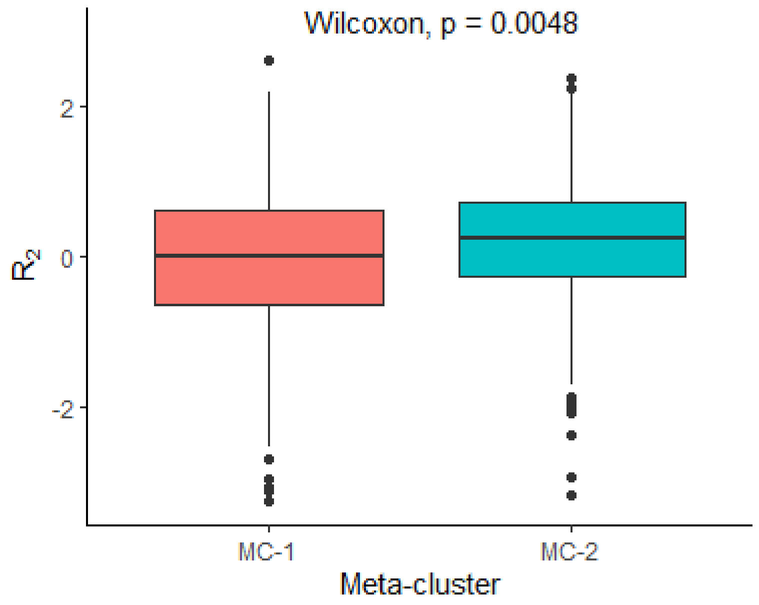


**Fig. S4. Box plot of MRI-derived frontal white matter $R_2$ by meta-cluster assignment.** The $R_2$ measure is derived from MRI data and is based on a voxel that captures the frontal white matter, as described in the methods. The MC-1 group has a lower median value, as illustrated in this box and whisker plot; the box is defined by measuring the lower quartile (25$^{th}$ percentile), median and upper quartile (75$^{th}$ percentile) of the values. The whiskers denote the two lines outside of the box that extend to the maximum and minimum values. *MC* meta-cluster.

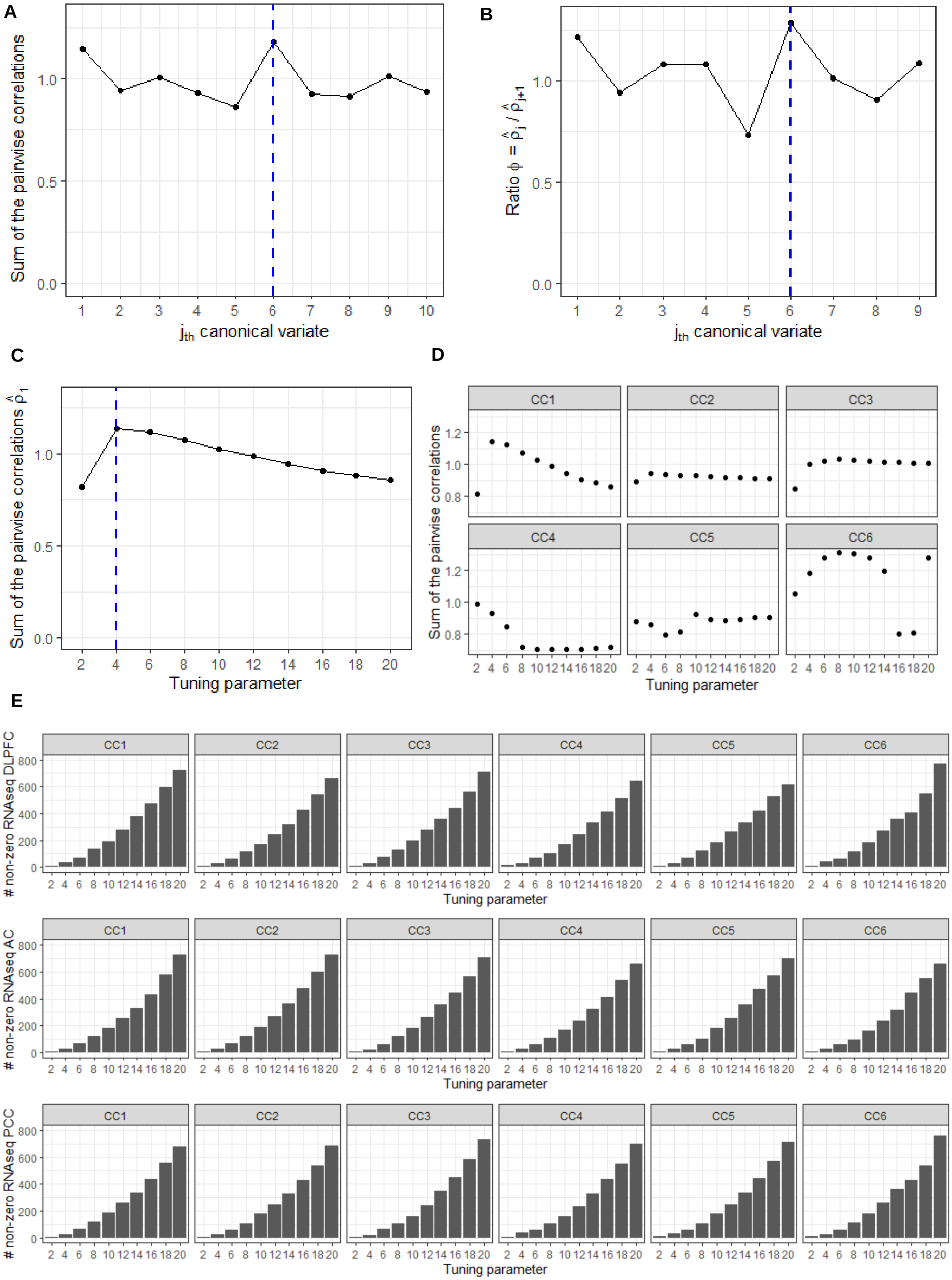
A
Sum of the pairwise correlations
jth canonical variate
B
Ratio ϕ = ρ̂j / ρ̂j+1
jth canonical variate
C
Sum of the pairwise correlations ρ̂1
Tuning parameter
D
CC1
CC2
CC3
CC4
CC5
CC6
Sum of the pairwise correlations
Tuning parameter
E
### non-zero RNAseq DLPFC
### non-zero RNAseq AC
### non-zero RNAseq PCC
Tuning parameter

**Fig. S5. Sum of the pairwise correlations from the three brain regions under various sparse parameters by canonical variates.** (**A**) Distribution of sum of the pairwise correlations between $j$th canonical variates from the three brain regions ($\hat{\rho}_j$), ranging from $j$=1 to 10. (**B**) Distribution of eigenvalues ratio $\hat{\varphi}_j = \hat{\rho}_j/\hat{\rho}_{j+1}$ where $j$=1 to 9. The ratio was maximized at $j$ =6 and first six canonical variates in each region were used for further analysis. (**C**) Distribution of sum of the pairwise correlations between first canonical variates from the three regions ($\hat{\rho}_1$) under various sparse parameters sets ranging from (2,2,2) to (20,20,20) where sparse parameter for each brain region ranged from 2 to 20 increasing by 2. (**D**) Distribution of sum of the pairwise correlations between $j$th canonical variates from the three brain regions $\hat{\rho}_j$ for $j$=1 to 6 using sparse parameters sets ranging from (2,2,2) to (20,20,20). Sparse parameters set of (4,4,4) had an overall large sum of the correlations and was selected for modeling. (**E**) Number of genes contributing to each canonical variate in three brain regions selected from deploying a sparse multiple CCA that maximize the sum of the pairwise correlations from the three brain regions under the various sparse parameters sets ranging from (2,2,2) to (20,20,20).

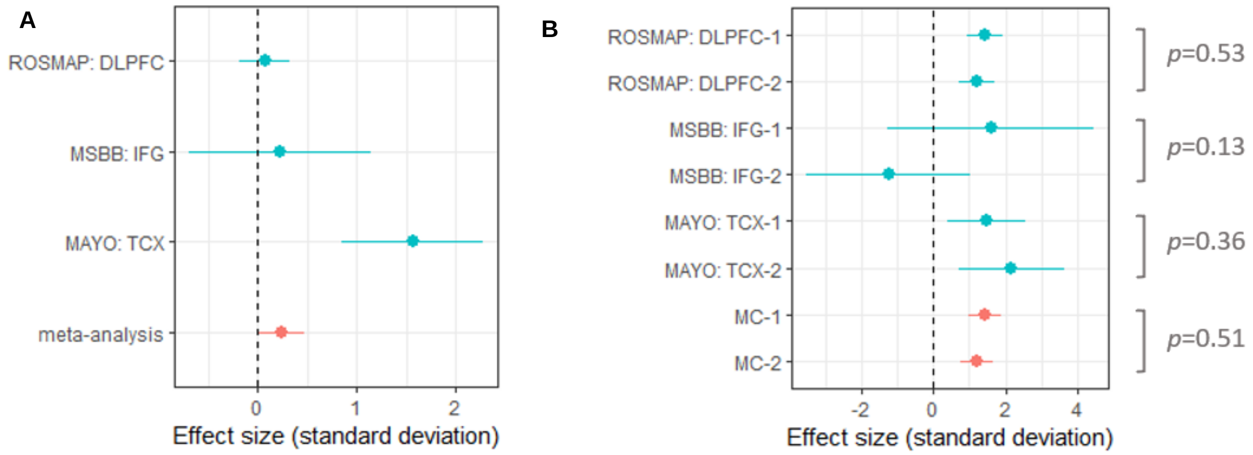


**Fig. S6. Validation study in Mount Sinai Brain Bank and Mayo Clinic.** (**A**) Association of pathologic diagnosis of AD or (**B**) interaction of the effect of *APOEε4* on the pathological AD between the two meta-clusters in ROSMAP, MSBB, and Mayo, after adjusting for age and sex. *p* p-value of testing the difference in the effect *APOEε4* on the pathological AD between the two meta-clusters, i.e., testing the interaction of MC with *APOEε4* on the pathological AD. *ROSMAP* Religious Order Study and Memory and Aging Project; *MSBB* Mount Sinai Brain Bank; *Mayo* Mayo Clinic; *DLPFC* dorsolateral prefrontal cortex; *IFG* inferior frontal gyrus; *TCX* temporal cortex; *meta-analysis* represents the meta-analysis of results from ROSMAP, MSBB, and Mayo (red line).

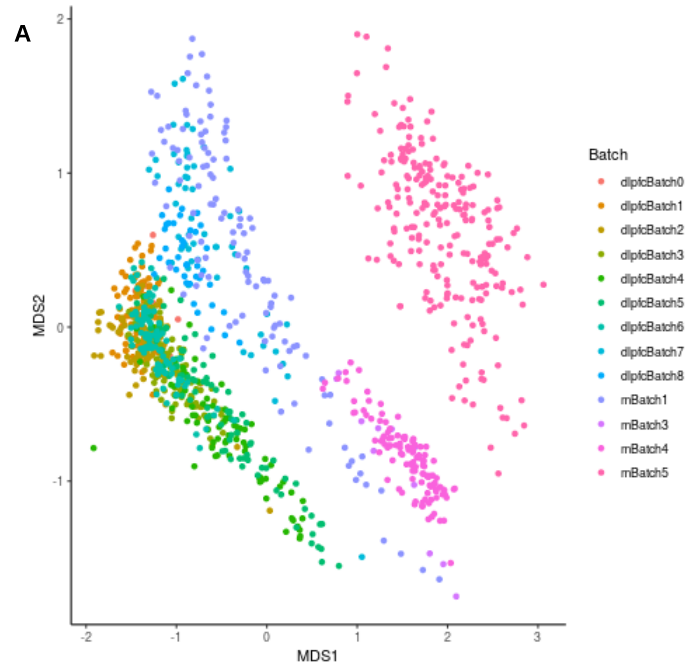


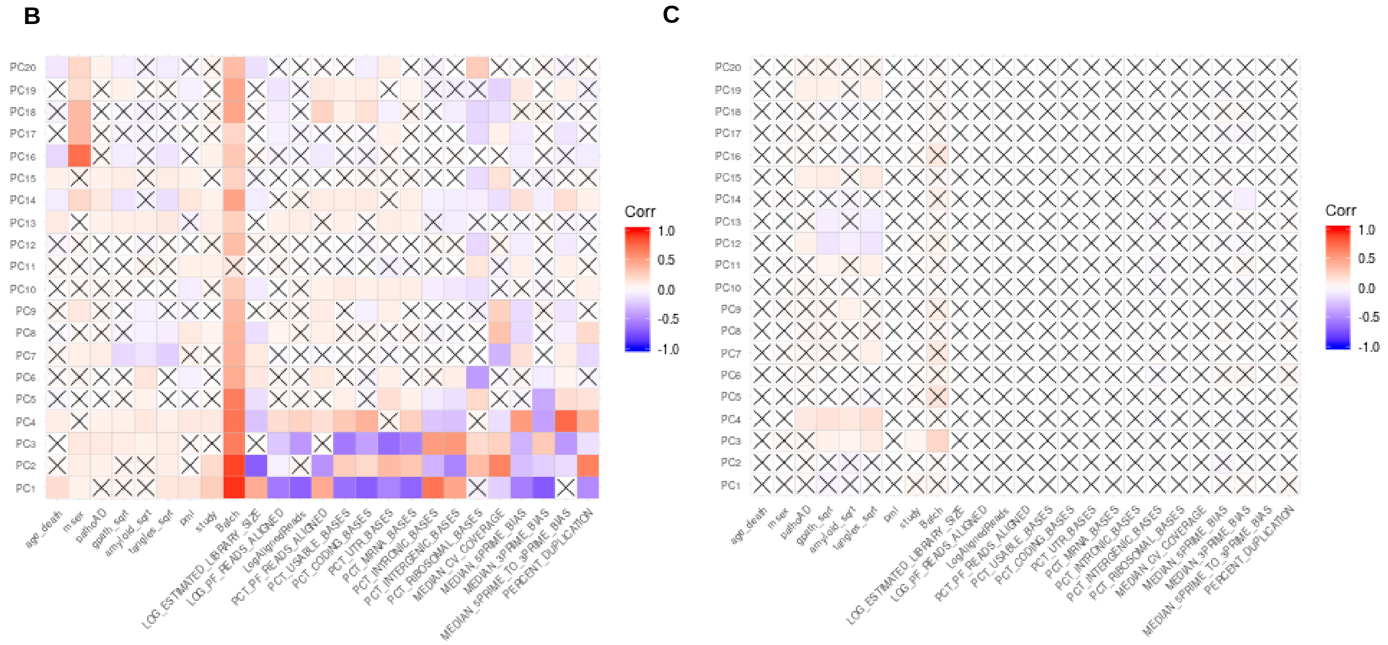


**Fig. S7. Exploring covariates to be adjusted for expression modeling.** (**A**) Distribution of participants by batch using MDS plot of normalized TPM values in DLPFC. (**B**) Correlation matrix between covariates and first 20 principal components from the normalized TPM values. (**C**) Correlation matrix between covariates and first 20 principal components from the residuals of the normalized TPM values (measure of gene expression after regressing out the differences between samples, studies, experimental batch effects or unwanted RNA-sequencing specific technical variations). X mark represents the correlation is not significant ($p>0.05$).

**Table S1. Distribution of participants by clusters obtained from each brain region and meta-clusters. (A)** Number of participants in Discovery and Replication cohorts by subgroups identified in each brain region and meta-clusters.

| | Discovery | | Replication | | |
|---|---|---|---|---|---|
| | 1 | 2 | 1 | 2 | Missing subjects |
| AC | 270 | 189 | 100 | 172 | 418 |
| DLPFC | 241 | 218 | 305 | 328 | 57 |
| PCC | 188 | 271 | 64 | 138 | 488 |
| MC | 274 | 185 | 344 | 346 | 0 |

*Discovery* participants with an RNA-seq profile in all 3 regions; *Replication* those participants with an RNA-seq profile in only one or two regions; *AC* anterior caudate; *DLPFC* dorsolateral prefrontal cortex; *PCC* posterior cingulate cortex; *MC* meta-cluster.

**(B)** Number of Discovery participants by meta-clusters with subgroups identified in each brain region simultaneously.

<table>
<tr><th colspan="6" rowspan="2">n</th><th colspan="2">AC</th></tr>
<tr><th>1</th><th>2</th></tr>
<tr><td rowspan="8">MC</td><td rowspan="4">1</td><td rowspan="4">PCC</td><td rowspan="2">1</td><td rowspan="2">DLPFC</td><td>1</td><td>84</td><td>15</td></tr>
<tr><td>2</td><td>67</td><td>0</td></tr>
<tr><td rowspan="2">2</td><td rowspan="2">DLPFC</td><td>1</td><td>19</td><td>0</td></tr>
<tr><td>2</td><td>0</td><td>0</td></tr>
<tr><td rowspan="4">2</td><td rowspan="4">PCC</td><td rowspan="2">1</td><td rowspan="2">DLPFC</td><td>1</td><td>0</td><td>0</td></tr>
<tr><td>2</td><td>0</td><td>22</td></tr>
<tr><td rowspan="2">2</td><td rowspan="2">DLPFC</td><td>1</td><td>0</td><td>100</td></tr>
<tr><td>2</td><td>19</td><td>133</td></tr>
</table>

*AC* anterior caudate; *DLPFC* dorsolateral prefrontal cortex; *PCC* posterior cingulate cortex; *MC* meta-cluster; *n* sample size.

**Table S2.** Association of traits with meta-cluster assignment, after adjusting for age and sex.

| | Discovery (n=459) | | | | Replication (n=690) | | | | Meta-Analysis (n=1,149) | | | |
|---|---|---|---|---|---|---|---|---|---|---|---|---|
| Outcome | b | SE | $p$ | $p_{FDR}$ | b | SE | $p$ | $p_{FDR}$ | b | SE | $p$ | $p_{FDR}$ |
| Cognitive decline | -0.032 | 0.009 | 0.0002 | 0.003 | -0.020 | 0.007 | 0.008 | 0.042 | -0.025 | 0.006 | $7.27 \times 10^{-6}$ | 0.0001 |
| AD dementia | 0.504 | 0.241 | 0.036 | 0.085 | 0.590 | 0.193 | 0.002 | 0.031 | 0.556 | 0.151 | 0.0002 | 0.002 |
| Residual cognition | -0.229 | 0.093 | 0.014 | 0.067 | -0.194 | 0.074 | 0.009 | 0.042 | -0.208 | 0.058 | 0.0003 | 0.002 |
| Amyloid | 0.252 | 0.109 | 0.021 | 0.074 | 0.185 | 0.084 | 0.028 | 0.098 | 0.210 | 0.067 | 0.002 | 0.006 |
| Pathologic AD | 0.389 | 0.198 | 0.050 | 0.100 | 0.337 | 0.169 | 0.046 | 0.128 | 0.359 | 0.129 | 0.005 | 0.015 |
| Neocortical Lewy bodies | 0.803 | 0.305 | 0.009 | 0.060 | 0.302 | 0.223 | 0.174 | 0.346 | 0.476 | 0.180 | 0.008 | 0.019 |
| Tau | 0.271 | 0.123 | 0.028 | 0.078 | 0.148 | 0.102 | 0.148 | 0.346 | 0.198 | 0.079 | 0.012 | 0.023 |
| TDP-43 | 0.134 | 0.221 | 0.545 | 0.693 | 0.227 | 0.177 | 0.198 | 0.346 | 0.191 | 0.138 | 0.166 | 0.276 |
| Hippocampal sclerosis | 0.465 | 0.347 | 0.179 | 0.314 | 0.178 | 0.267 | 0.504 | 0.642 | 0.285 | 0.212 | 0.178 | 0.276 |
| Arteriolosclerosis | 0.137 | 0.201 | 0.494 | 0.692 | -0.178 | 0.162 | 0.270 | 0.420 | -0.054 | 0.126 | 0.667 | 0.841 |
| Cerebral atherosclerosis | 0.008 | 0.201 | 0.968 | 0.968 | -0.093 | 0.164 | 0.572 | 0.667 | -0.053 | 0.127 | 0.680 | 0.841 |
| MF PAM | -0.004 | 0.016 | 0.825 | 0.889 | -0.002 | 0.012 | 0.840 | 0.904 | -0.003 | 0.010 | 0.767 | 0.841 |
| IT PAM | 0.008 | 0.014 | 0.603 | 0.703 | -0.001 | 0.011 | 0.952 | 0.952 | 0.003 | 0.009 | 0.786 | 0.841 |
| Cerebral amyloid angiopathy | 0.189 | 0.208 | 0.363 | 0.565 | -0.129 | 0.160 | 0.420 | 0.587 | -0.011 | 0.127 | 0.931 | 0.931 |

The MC-2 is the reference group. *MC* meta-cluster; *Discovery* participants with an RNA-seq profile in all 3 regions; *Replication* participants with an RNA-seq profile in only one or two regions; *MF* Midfrontal cortex; *IT* Inferior temporal cortex; *PAM* the proportion of active microglia; *n* sample size; *b* estimate; *SE* standard error; $p$ p-value; $p_{FDR}$ FDR adjusted p-value using the tests in the table.

**Table S3**. Meta-analysis association of cognitive decline between MC-1 and MC-2, after adjusting for covariate(s) in addition to age and sex.

| Adjusted covariate(s) | n | b | SE | $p$ | $p_{FDR}$ |
|---|---|---|---|---|---|
| Amyloid | 1069 | -0.022 | 0.005 | $5.56\times 10^{-5}$ | $1.25\times 10^{-4}$ |
| Tau | 1073 | -0.021 | 0.005 | $1.08\times 10^{-5}$ | $4.86\times 10^{-5}$ |
| Amyloid and tau | 1068 | -0.020 | 0.005 | $3.48\times 10^{-5}$ | $1.04\times 10^{-4}$ |
| Pathologic AD | 1081 | -0.021 | 0.005 | $7.53\times 10^{-5}$ | $1.32\times 10^{-4}$ |
| AD dementia | 740 | -0.007 | 0.005 | 0.164 | 0.164 |
| $R_2$ | 532 | -0.028 | 0.008 | $2.24\times 10^{-4}$ | $2.88\times 10^{-4}$ |
| $R_2$, amyloid and tau | 527 | -0.023 | 0.007 | $3.65\times 10^{-4}$ | $4.10\times 10^{-4}$ |
| *CLU* | 841 | -0.029 | 0.006 | $6.91\times 10^{-6}$ | $4.86\times 10^{-5}$ |
| Braak stage | 1081 | -0.020 | 0.005 | $8.83\times 10^{-5}$ | $1.32\times 10^{-4}$ |

The MC-2 is the reference group. *MC* meta-cluster; *Adjusted covariates* adjusted pathologic indices; *n* sample size; *b* estimate; *SE* standard error; *p* p-value; $p_{FDR}$ FDR adjusted p-value using the tests in the table.

**Table S4.** Meta-analysis of association of other factors between MC-1 and MC-2, after adjusting for age and sex.

| Category | Outcome | n | b | SE | *p* |
|---|---|---|---|---|---|
| Epigenomic tau score | H3K9ac score | 520 | 16.329 | 3.343 | $1.04\times 10^{-6}$ |
| | H3K9ac score, adjusting for tau | 277 | 15.389 | 3.222 | $1.79\times 10^{-6}$ |
| $R_2$ | $R_2$ | 557 | -0.217 | 0.081 | 0.007 |
| Genetic load for AD | *APOEε4* | 1138 | 0.255 | 0.138 | 0.064 |
| | *APOEε2* | 1138 | -0.548 | 0.175 | 0.002 |
| *APOE* genotype | *ε2/ε2* vs *ε3/ε3* | 1138 | -13.793 | 303.685 | 0.964 |
| | *ε2/ε3* vs *ε3/ε3* | 1138 | -0.476 | 0.193 | 0.014 |
| | *ε2/ε4* vs *ε3/ε3* | 1138 | -0.279 | 0.434 | 0.520 |
| | *ε3/ε4* vs *ε3/ε3* | 1138 | 0.148 | 0.149 | 0.322 |
| | *ε4/ε4* vs *ε3/ε3* | 1138 | 1.089 | 0.535 | 0.042 |
| Polygenic risk score for clinical diagnosis of AD | IGAP | 860 | 0.075 | 0.069 | 0.277 |
| | IGAP without APOE/TOMM40 | 860 | 0.062 | 0.069 | 0.363 |
| | CCACE | 860 | 0.032 | 0.067 | 0.637 |
| | CCACE without APOE/TOMM40 | 860 | 0.014 | 0.067 | 0.840 |
| Cell type proportion | Neurons | 1092 | -0.063 | 0.005 | $1.12\times 10^{-37}$ |
| | Neurons, adjusting for *SNAP25* | 912 | -0.057 | 0.005 | $5.06\times 10^{-28}$ |
| | Astrocytes | 1092 | 0.027 | 0.003 | $8.04\times 10^{-20}$ |
| | Oligodendrocytes | 1092 | 0.012 | 0.002 | $4.98\times 10^{-8}$ |
| | Microglia | 1092 | 0.008 | 0.001 | $6.82\times 10^{-16}$ |
| | Endothelial cells | 1092 | 0.021 | 0.002 | $5.03\times 10^{-17}$ |
| Cell type proportion for cognitively non-impaired subjects | Neurons | 204 | -0.048 | 0.010 | $2.62\times 10^{-6}$ |
| | Astrocytes | 204 | 0.030 | 0.007 | $1.22\times 10^{-5}$ |
| | Oligodendrocytes | 204 | 0.005 | 0.005 | 0.272 |
| | Microglia | 204 | 0.012 | 0.002 | $2.29\times 10^{-7}$ |
| | Endothelial cells | 204 | 0.016 | 0.006 | 0.009 |
| *SNAP25* | *SNAP25* | 912 | -0.076 | 0.015 | $3.80\times 10^{-7}$ |
| | *SNAP25*, adjusting for neurons | 912 | -0.056 | 0.016 | $6.38\times 10^{-4}$ |
| *CLU* | *CLU* | 898 | 0.126 | 0.047 | 0.007 |
| Aged microglia gene set | HuMi-Aged | 1048 | 0.282 | 0.025 | $1.84\times 10^{-30}$ |
| Epigenomic Factor of Activated Microglia | EFAM | 456 | 0.224 | 0.131 | 0.087 |
| Braak stage | I | 1149 | 0.454 | 0.899 | 0.613 |
| | II | 1149 | 1.011 | 0.876 | 0.248 |
| | III | 1149 | 0.882 | 0.855 | 0.302 |
| | IV | 1149 | 0.924 | 0.851 | 0.277 |
| | V | 1149 | 1.216 | 0.854 | 0.155 |
| | VI | 1149 | 0.969 | 1.052 | 0.357 |
| CERAD score | Definite AD | 1149 | 0.490 | 0.167 | 0.003 |

| | | | | | |
|---|---|---|---|---|---|
| | Probable AD | 1149 | 0.352 | 0.162 | 0.029 |
| | Possible AD | 1149 | 0.227 | 0.238 | 0.341 |

The MC-2 is the reference group. *MC* meta-cluster; *Outcome* other factors; *n* sample size; *b* estimate; *SE* standard error; *p* p-value.

**Table S5**. Meta-analysis of association of factor between MC-1 and MC-2 among participants with pathological AD, among those without pathological AD, among those with AD dementia, or among those without AD dementia, after adjusting for age and sex.

| | With pathological AD | | | | Without pathological AD | | | | With AD dementia | | | | Without AD dementia | | | |
|---|---|---|---|---|---|---|---|---|---|---|---|---|---|---|---|---|
| Factor | b | SE | $p$ | $p_{FDR}$ | b | SE | $p$ | $p_{FDR}$ | b | SE | $p$ | $p_{FDR}$ | b | SE | $p$ | $p_{FDR}$ |
| Amyloid | 0.100 | 0.061 | 0.100 | 0.203 | 0.024 | 0.072 | 0.742 | 0.912 | 0.106 | 0.105 | 0.311 | 0.686 | 0.045 | 0.111 | 0.682 | 0.919 |
| Tau | 0.114 | 0.102 | 0.263 | 0.361 | 0.044 | 0.074 | 0.552 | 0.912 | 0.020 | 0.151 | 0.897 | 0.898 | 0.014 | 0.088 | 0.873 | 0.923 |
| Cognitive decline | -0.025 | 0.007 | 0.001 | 0.004 | -0.014 | 0.007 | 0.045 | 0.225 | -0.002 | 0.009 | 0.795 | 0.898 | -0.007 | 0.004 | 0.043 | 0.213 |
| Cognitive decline, adjusting for amyloid and tau | -0.023 | 0.006 | 0.0002 | 0.003 | -0.014 | 0.007 | 0.045 | 0.225 | -0.006 | 0.008 | 0.461 | 0.692 | -0.007 | 0.004 | 0.042 | 0.213 |
| Cognitive decline, adjusting for amyloid,tau, and $R_2$ | -0.026 | 0.008 | 0.001 | 0.005 | -0.019 | 0.011 | 0.077 | 0.288 | -0.011 | 0.011 | 0.324 | 0.686 | -0.011 | 0.005 | 0.023 | 0.213 |
| Residual cognition | -0.254 | 0.076 | 0.001 | 0.004 | -0.130 | 0.089 | 0.143 | 0.358 | -0.017 | 0.097 | 0.858 | 0.898 | -0.033 | 0.058 | 0.567 | 0.851 |
| TDP-43 | 0.092 | 0.162 | 0.570 | 0.610 | 0.315 | 0.279 | 0.258 | 0.553 | -0.119 | 0.215 | 0.582 | 0.793 | -0.100 | 0.323 | 0.756 | 0.919 |
| Cerebral atherosclerosis | -0.131 | 0.161 | 0.418 | 0.522 | 0.144 | 0.214 | 0.500 | 0.912 | -0.183 | 0.214 | 0.391 | 0.686 | 0.224 | 0.237 | 0.346 | 0.728 |
| Cerebral amyloid angiopathy | -0.094 | 0.151 | 0.533 | 0.610 | -0.129 | 0.262 | 0.621 | 0.912 | -0.026 | 0.207 | 0.898 | 0.898 | 0.023 | 0.234 | 0.923 | 0.923 |
| Arteriolosclerosis | -0.047 | 0.159 | 0.770 | 0.770 | -0.049 | 0.210 | 0.815 | 0.912 | -0.171 | 0.209 | 0.412 | 0.686 | 0.289 | 0.238 | 0.225 | 0.676 |
| Hippocampal sclerosis | 0.387 | 0.250 | 0.122 | 0.204 | -0.021 | 0.471 | 0.964 | 0.964 | -0.308 | 0.263 | 0.241 | 0.686 | 0.989 | 1.272 | 0.437 | 0.728 |
| Neocortical Lewy bodies | 0.382 | 0.214 | 0.074 | 0.184 | 0.550 | 0.346 | 0.112 | 0.336 | 0.284 | 0.257 | 0.270 | 0.686 | -0.119 | 0.463 | 0.797 | 0.919 |

| | | | | | | | | | | | | | | | | |
|---|---|---|---|---|---|---|---|---|---|---|---|---|---|---|---|---|
| *APOEε4* | 0.182 | 0.163 | 0.265 | 0.361 | 0.111 | 0.296 | 0.708 | 0.912 | 0.344 | 0.217 | 0.112 | 0.561 | -0.282 | 0.288 | 0.328 | 0.728 |
| *APOEε2* | -0.382 | 0.238 | 0.108 | 0.203 | -0.626 | 0.267 | 0.019 | 0.225 | -0.748 | 0.313 | 0.017 | 0.147 | -0.403 | 0.279 | 0.148 | 0.555 |
| $R_2$ | -0.280 | 0.098 | 0.004 | 0.012 | 0.027 | 0.144 | 0.851 | 0.912 | -0.326 | 0.140 | 0.020 | 0.147 | -0.102 | 0.123 | 0.408 | 0.728 |

The MC-2 is the reference group. *MC* meta-cluster; *b* estimate; *SE* standard error; *p* p-value;

$p_{FDR}$ FDR adjusted p-value using the tests in the table.

**Table S6**. Meta-analysis association of AD-related gene between MC-1 and MC-2, after adjusting for age and sex.

| Gene | SNP | Chr | bp | Minor allele | Major allele | Minor freq | b | SE | *p* | $p_{FDR}$ |
|---|---|---|---|---|---|---|---|---|---|---|
| *CLU* | rs9331896 | 8 | 27467686 | C | T | 0.407 | 0.126 | 0.047 | 0.007 | 0.129 |
| *CR1* | rs4844610 | 1 | 207802552 | A | C | 0.194 | -0.067 | 0.038 | 0.080 | 0.443 |
| *CD2AP* | rs9473117 | 6 | 47431284 | C | A | 0.271 | -0.064 | 0.042 | 0.127 | 0.443 |
| *BIN1* | rs6733839 | 2 | 127892810 | T | C | 0.394 | 0.067 | 0.045 | 0.134 | 0.443 |
| *CASS4* | rs6024870 | 20 | 54997568 | A | G | 0.079 | -0.039 | 0.026 | 0.134 | 0.443 |
| *FERMT2* | rs17125924 | 14 | 53391680 | G | A | 0.091 | 0.035 | 0.027 | 0.208 | 0.443 |
| *NYAP1g* | rs12539172 | 7 | 100091795 | T | C | 0.317 | 0.055 | 0.044 | 0.217 | 0.443 |
| *ECHDC3* | rs7920721 | 10 | 11720308 | G | A | 0.37 | 0.049 | 0.046 | 0.280 | 0.443 |
| *PICALM* | rs3851179 | 11 | 85868640 | T | C | 0.358 | -0.049 | 0.046 | 0.282 | 0.443 |
| *SPI1h* | rs3740688 | 11 | 47380340 | G | T | 0.482 | -0.051 | 0.049 | 0.302 | 0.443 |
| *ABCA7* | rs3752246 | 19 | 1056492 | G | C | 0.184 | -0.037 | 0.037 | 0.312 | 0.443 |
| *ACE* | rs138190086 | 17 | 61538148 | A | G | 0.015 | 0.011 | 0.011 | 0.322 | 0.443 |
| *EPHA1* | rs10808026 | 7 | 143099133 | A | C | 0.202 | 0.037 | 0.037 | 0.323 | 0.443 |
| *INPP5D* | rs10933431 | 2 | 233981912 | G | C | 0.233 | -0.039 | 0.040 | 0.326 | 0.443 |
| *SLC24A4* | rs12881735 | 14 | 92932828 | C | T | 0.229 | -0.033 | 0.040 | 0.416 | 0.527 |
| *HLA_DRB1* | rs9271058 | 6 | 32575406 | A | T | 0.292 | -0.022 | 0.044 | 0.615 | 0.717 |
| *PTK2B* | rs73223431 | 8 | 27219987 | T | C | 0.347 | -0.021 | 0.045 | 0.642 | 0.717 |
| *SORL1* | rs11218343 | 11 | 121435587 | C | T | 0.031 | 0.006 | 0.016 | 0.710 | 0.728 |
| *MS4A2* | rs7933202 | 11 | 59936926 | C | A | 0.414 | 0.017 | 0.048 | 0.728 | 0.728 |

The MC-2 is the reference group. *MC* meta-cluster; *SNP* single nucleotide polymorphism; *Chr* chromosome; *bp* base pairs; *Minor allele* Minor/Risk allele; *freq* frequency; *b* estimate; *SE* standard error; *p* p-value; $p_{FDR}$ FDR adjusted p-value using the tests in the table.

**Table S7.** Strength of evidence for interaction of meta-cluster assignment with predictor on outcome, after adjusting for age and sex.

| Outcome | Predictor | MC | n | b | SE | $p$ | $p^*$ |
|---|---|---|---|---|---|---|---|
| Cognitive decline | *APOEε4* | 1 | 499 | -0.0686 | 0.0091 | $5.63\times 10^{-14}$ | 0.003 |
| | | 2 | 578 | -0.0322 | 0.0085 | $1.44\times 10^{-4}$ | |
| | *APOE* | 1 | 499 | 0.0248 | 0.0145 | 0.087 | 0.190 |
| | | 2 | 578 | -0.0281 | 0.0377 | 0.455 | |
| | *APOEε4*, adjusting for amyloid | 1 | 493 | -0.0505 | 0.0096 | $1.29\times 10^{-7}$ | 0.006 |
| | | 2 | 572 | -0.0156 | 0.0085 | 0.067 | |
| | *APOEε4*, adjusting for tau | 1 | 495 | -0.0336 | 0.0084 | $7.07\times 10^{-5}$ | 0.029 |
| | | 2 | 574 | -0.0090 | 0.0074 | 0.224 | |
| | *APOEε4*, adjusting for amyloid and tau | 1 | 492 | -0.0304 | 0.0088 | $5.46\times 10^{-4}$ | 0.037 |
| | | 2 | 572 | -0.0060 | 0.0076 | 0.428 | |
| | *APOEε4*, adjusting for EFAM | 1 | 206 | -0.0759 | 0.0149 | $3.62\times 10^{-7}$ | $9.37\times 10^{-4}$ |
| | | 2 | 223 | -0.0125 | 0.0120 | 0.301 | |
| | *APOEε4*, adjusting for HuMi-Aged | 1 | 444 | -0.0656 | 0.0098 | $2.08\times 10^{-11}$ | 0.010 |
| | | 2 | 541 | -0.0316 | 0.0088 | $3.41\times 10^{-4}$ | |
| | *APOEε4*, adjusting for *SNAP25* | 1 | 390 | -0.0624 | 0.0105 | $3.19\times 10^{-9}$ | 0.015 |
| | | 2 | 470 | -0.0281 | 0.0094 | $2.73\times 10^{-3}$ | |
| | $R_2$ | 1 | 253 | 0.0299 | 0.0060 | $5.48\times 10^{-7}$ | 0.253 |
| | | 2 | 279 | 0.0205 | 0.0056 | $2.63\times 10^{-4}$ | |
| | Epigenomic perturbation score (H3K9ac_score) | 1 | 222 | -0.0005 | 0.0002 | 0.015 | 0.297 |
| | | 2 | 269 | -0.0002 | 0.0001 | 0.123 | |
| | *SNAP25* | 1 | 392 | 0.0472 | 0.0187 | 0.012 | 0.057 |
| | | 2 | 471 | 0.0995 | 0.0201 | $7.68\times 10^{-7}$ | |
| | Epigenomic factor of activated microglia (EFAM) | 1 | 206 | -0.0195 | 0.0050 | $1.05\times 10^{-4}$ | 0.029 |
| | | 2 | 223 | -0.0055 | 0.0040 | 0.175 | |
| Tau | *APOEε4* | 1 | 517 | 0.9563 | 0.1315 | $3.50\times 10^{-13}$ | 0.037 |
| | | 2 | 613 | 0.5903 | 0.1164 | $3.91\times 10^{-7}$ | |
| | Amyloid | 1 | 522 | 0.5797 | 0.0486 | $7.23\times 10^{-33}$ | 0.024 |
| | | 2 | 614 | 0.4364 | 0.0412 | $3.69\times 10^{-26}$ | |

The MC-2 is the reference group. *MC* meta-cluster; *n* sample size; *b* estimate; *SE* standard error; *p* p-value of testing the effect of predictor on outcome within each MC; $p^*$ p-value of testing the difference in the effect of predictor on outcome between MC-1 and MC-2, i.e., testing the interaction of MC with the predictor on outcome.

**Table S8**. Strength of evidence for interaction of meta-cluster assignment with *APOEε4* on cognitive decline among participants with pathological AD, among those without pathological AD, among those with AD dementia, or among those without AD dementia, after adjusting for age and sex.

| Outcome | Predictor | Participants | MC | n | b | SE | *p* | $p^*$ |
|---|---|---|---|---|---|---|---|---|
| Cognitive decline | *APOEε4* | With pathological AD | 1 | 336 | -0.065 | 0.011 | $4.46 \times 10^{-9}$ | 0.005 |
| | | | 2 | 348 | -0.021 | 0.011 | 0.056 | |
| | | Without pathological AD | 1 | 163 | -0.019 | 0.017 | 0.245 | 0.767 |
| | | | 2 | 230 | -0.013 | 0.013 | 0.317 | |
| | | With AD dementia | 1 | 209 | -0.042 | 0.012 | 0.0004 | 0.113 |
| | | | 2 | 172 | -0.012 | 0.015 | 0.419 | |
| | | Without AD dementia | 1 | 148 | -0.006 | 0.008 | 0.463 | 0.768 |
| | | | 2 | 208 | -0.003 | 0.006 | 0.632 | |

The MC-2 is the reference group. *MC* meta-cluster; *n* sample size; *b* estimate; *SE* standard error; *p* p-value of testing the effect of predictor on outcome within each MC; $p^*$ p-value of testing the difference in the effect of predictor on outcome between MC-1 and MC-2, i.e., testing the interaction of MC with the predictor on outcome.

**Table S9**. Estimated cumulative risk of AD dementia onset in MC-1 and MC-2, adjusting for sex and education in ROSMAP ($\times 10^{-2}$).

| | MC-1 | | | MC-2 | | |
|---|---|---|---|---|---|---|
| Age | Risk | Lower limit | Upper limit | Risk | Lower limit | Upper limit |
| 70 | 9.63 | 1.71 | 22.04 | 6.63 | 1.46 | 14.89 |
| 75 | 16.29 | 5.17 | 30.07 | 11.63 | 4.17 | 21.44 |
| 80 | 26.92 | 13.41 | 40.80 | 20.01 | 10.46 | 30.78 |
| 85 | 41.85 | 28.11 | 54.26 | 32.57 | 21.92 | 43.17 |
| 90 | 60.57 | 48.96 | 70.37 | 49.80 | 39.32 | 59.17 |
| 95 | 79.39 | 70.63 | 86.13 | 69.60 | 60.24 | 77.15 |
| 100 | 93.13 | 87.23 | 96.72 | 87.21 | 79.60 | 92.56 |

*MC* meta-cluster;

**Table S10**. Meta-analysis association of pathologic diagnosis of AD between MC-1 and MC-2 in ROSMAP, MSBB and Mayo, after adjusting for age and sex.

| Trait | Cohort | n | b | SE | p |
|---|---|---|---|---|---|
| Pathologic AD | ROSMAP: DLPFC | 1092 | 0.070 | 0.130 | 0.591 |
| | MSBB: IFG | 186 | 0.230 | 0.468 | 0.623 |
| | Mayo: TCX | 262 | 1.565 | 0.365 | $1.83\times 10^{-5}$ |
| | meta-analysis | 1540 | 0.237 | 0.118 | 0.045 |

The MC-2 is the reference group. *MC* meta-cluster; *n* sample size; *b* estimate; *SE* standard error; *p* p-value. *ROSMAP* Religious Order Study and Memory and Aging Project; *MSBB* Mount Sinai Brain Bank; *Mayo* Mayo Clinic; *DLPFC* dorsolateral prefrontal cortex; *IFG* inferior frontal gyrus; *TCX* temporal cortex; *meta-analysis* represents the meta-analysis of results from ROSMAP, MSBB, and Mayo.

**Table S11**. Strength of evidence for interaction of meta-cluster assignment with *APOEε4* on pathologic diagnosis of AD, in ROSMAP, MSBB and Mayo, after adjusting for age and sex.

| Outcome | Predictor | Cohort | MC | n | b | SE | *p* | $p^*$ |
|---|---|---|---|---|---|---|---|---|
| Pathologic AD | *APOEε4* | ROSMAP: DLPFC | 1 | 516 | 1.425 | 0.253 | $1.81\times 10^{-8}$ | |
| | | | 2 | 566 | 1.202 | 0.245 | $8.98\times 10^{-7}$ | 0.526 |
| | | MSBB: IFG | 1 | 37 | 1.591 | 1.474 | 0.281 | |
| | | | 2 | 28 | -1.250 | 1.166 | 0.284 | 0.131 |
| | | Mayo: TCX | 1 | 86 | 1.480 | 0.548 | 0.007 | |
| | | | 2 | 65 | 2.169 | 0.751 | 0.004 | 0.458 |
| | | meta-analysis | 1 | 639 | 1.439 | 0.227 | $2.37\times 10^{-10}$ | |
| | | | 2 | 659 | 1.198 | 0.228 | $1.53\times 10^{-7}$ | 0.454 |

The MC-2 is the reference group. *MC* meta-cluster; *n* sample size; *b* estimate; *SE* standard error; *p* p-value of testing the effect of predictor on outcome within each MC; $p^*$ p-value of testing the difference in the effect of *APOEε4* on pathologic diagnosis of AD between MC-1 and MC-2, i.e., testing the interaction of MC with the *APOEε4* on pathologic AD; *ROSMAP* Religious Order Study and Memory and Aging Project; *MSBB* Mount Sinai Brain Bank; *Mayo* Mayo Clinic; *DLPFC* dorsolateral prefrontal cortex; *IFG* inferior frontal gyrus; *TCX* temporal cortex; *meta-analysis* represents the meta-analysis of results from ROSMAP, MSBB, and Mayo.
